# Structural adaptations for enhanced translation kinetics in evolved ribosomes

**DOI:** 10.64898/2026.03.05.706023

**Authors:** Tushar Raskar, Alan Costello, Ahmed H. Badran, James S. Fraser

## Abstract

The ribosomal RNA sequence governs translation dynamics, yet understanding how changes beyond the conserved catalytic centers influence kinetics and protein yield remains limited. Using orthogonal ribosome phage-assisted continuous evolution (oRibo-PACE), we recently reported chimeric ribosomes derived from *Escherichia coli*, *Pseudomonas aeruginosa*, and *Vibrio cholerae* endowed with elevated orthogonal translation activity as compared to their starting counterparts. Here, we structurally characterize these kinetically enhanced ribosomes using cryo-electron microscopy and uncover a potential relationship between 16S rRNA stability and translation efficiency. Compared to their naive starting points, evolved ribosomes exhibit extensive RNA structural adaptation, often introduced by mismatches at key helical junctions, which leads to local RNA-protein rearrangements and destabilizes non-canonical base pairs. Compensatory mutations that restore base-pairing stability and eliminate flexibility reduced translational activity to wild-type levels. Across trajectories, increased translational output correlates with subtle, localized changes in the 16S rRNA sequence that introduce limited structural destabilization at specific elements. Taken together, our work provides new insights into rRNA structural malleability and establishes principles for engineering ribosomes with altered translation properties.

**Graphical abstract:** 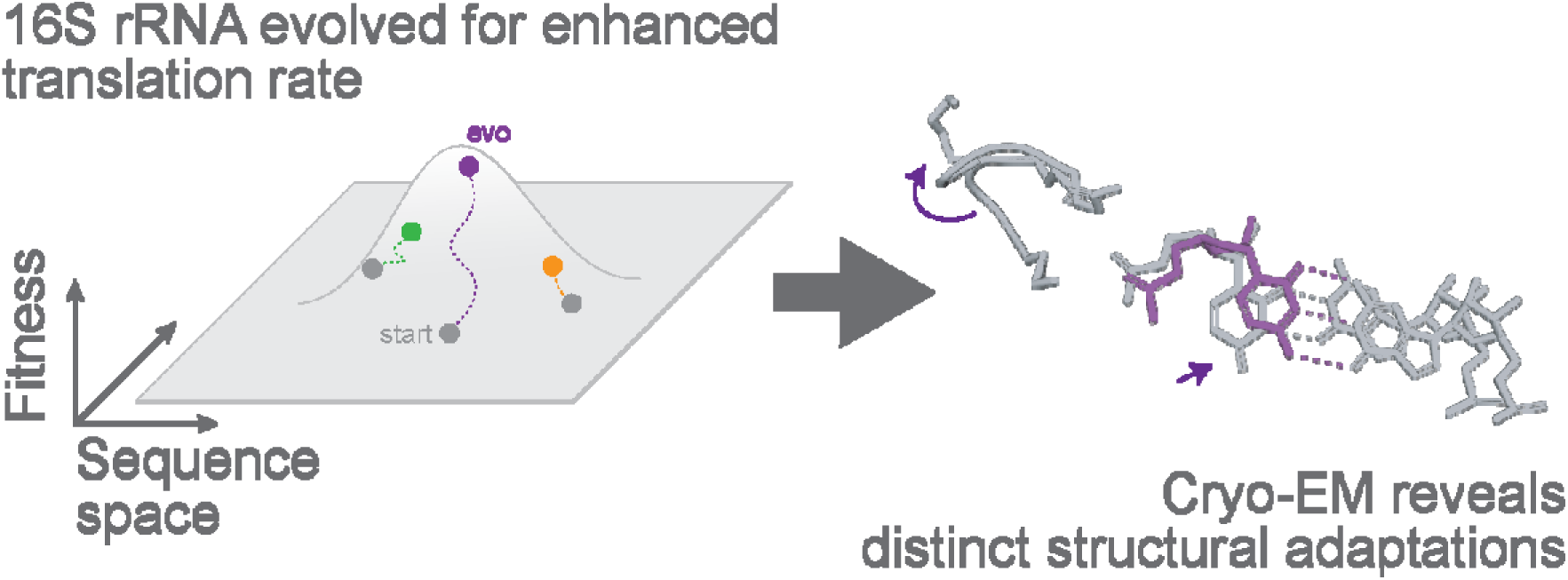

## Introduction

Ribosomes are highly conserved molecular machines responsible for synthesizing all cellular proteins by decoding genetic information and polymerizing amino acids into defined sequences. Latent functional capacity of the ribosome has been observed through programmed synthesis of polymers with diverse non-canonical building blocks(1–3), and decoding of alternative genetic codes(4–6). Unfortunately, many of these exciting non-standard polymerizations are low yielding and/or currently limited to in vitro translation systems. The ability to evolve new ribosome function has far-reaching applications, however, engineering essential biological components, such as the ribosome for these new-to-nature activities can be confounded by negative impacts on growth and survival.

Several attempts to orthogonalize the *E. coli* ribosome have been described to date, with successful decoupling of the ribosome 16S rRNA coming from creation of discrete transcription-translation networks(7, 8). These orthogonal translation systems (OTS) allow researchers to probe ribosome sequence-function relationships without perturbing host cell function. Directed evolution is a powerful approach for endowing biomolecules with novel functions(9, 10), and has recently been applied to OTSs for the discovery of ribosomes with augmented characteristics (7, 11, 12). Many of these studies have focused on the development of new-to-nature bioactivities, including polymerization of non-proteinogenic monomers and decoding of expanded quadruplet codons(13). Despite extensive research on ribosome evolution and engineering(14, 15), our understanding of regulatory sites beyond the well-characterized peptidyl transferase center and decoding regions remains limited(16). To date, few studies have explored the fundamental structure-function relationships of ribosome variants and how they relate to altered or completely novel activities.

We recently reported a technology to evolve ribosomal 16S rRNA for enhanced translation kinetics(17). Specifically, we combined ribosome engineering(18), 16S rRNA-mRNA orthogonalization(7), and phage-assisted continuous evolution (PACE)(18) to yield orthogonal ribosome PACE (oRibo-PACE), a system that enables researchers to readily tune the functions of heterologously expressed ribosomes in the workhorse bacterium *E. coli*.

While these chimeric evolved ribosomes showed enhanced orthogonal translation activity without concomitant reduction in fidelity, growth, or survival, the biochemical and/or biophysical bases for these phenotypes remained unknown. We hypothesized that these functional outcomes may be accompanied by specific structural changes since many mutations were observed in highly conserved regions of the 16S rRNA.

Here, we use single-particle cryo-electron microscopy (cryo-EM) to investigate the relationship between the observed biochemical adaptations and possible structural changes incurred during oRibo-PACE evolution. Our analyses integrated (1) structural differences between evolved 16S rRNA and their progenitor sequences, (2) conformational adaptations of *E. coli* r-proteins to heterologous rRNAs, and (3) the conformational changes between wild-type, chimeric, and evolved chimeric structures. These efforts revealed structural adaptations in *E. coli* r-proteins to accommodate heterologous 16S rRNAs(19), and identified structural changes that we posit may contribute to their enhanced translational capabilities. Taken together, this study provides new insights into how subtle structural adaptations can alter ribosome functions and endow them with novel properties.

## Methods

### Cell culture and scale up

The chimeric 16S strains for *P. aeruginosa*, *V. cholerae* and *E. coli* were inoculated in 12 mL Luria Bertani (LB) broth overnight. 1 mL of the starter culture was then added to 1000 mL LB broth and incubated at 37℃ at 180 rpm till the optical density (OD_600_) of the cells reaches 0.5-0.6. The cells were then harvested by centrifugation at 4000 g in a Beckman Coulter 8.1 rotor at 4℃ for 40 minutes. The media was then poured off followed by addition of 5 mL of pellet wash buffer (20 mM HEPES-KOH pH 7.5, 100 mM NaCl, 10 mM MgCl_2_) per pellet. The cells were homogenized in the pellet wash buffer followed by centrifugation at 4℃ at 4000 rpm for 20 minutes. The supernatant was removed and the pellet was stored at -80℃ until further use.

### Cell lysis and 70S ribosome purification

The cells were resuspended in lysis buffer (20 mM HEPES-KOH pH 7.5, 200 mM NH_4_Cl, 20 mM Mg(OAc)₂, 0.5 mM EDTA, 6 mM Beta-mercaptoethanol (β-ME), 10 U/mL SUPERase-In RNAse inhibitor, 0.1 mM PMSF) until homogenized followed by lysis using Emulsiflex C3 high pressure homogenizer. The lysate was then spun at 30,000 g for 40 minutes at 4℃. The supernatant was then sterile filtered through a 0.45 µ filter. 6 mL of filtered and cleared supernatant was layered on top 6 mL of 32% sucrose buffer (20 mM HEPES-KOH pH 7.5, 500 mM NH_4_Cl, 20 mM Mg(OAc)_2_, 0.5 mM EDTA, 6 mM β-ME, 10 U/mL SUPERase-In inhibitor) in Seton #7030 Ultracentrifuge Tubes. The tubes were then centrifuged at 35,000 rpm at 4℃ for 4 hours in the SW41Ti rotor. The supernatant was discarded and the transparent pellet was resuspended in buffer A (20 mM HEPES-KOH pH 7.5, 200 mM NH₄Cl, 20 mM Mg(OAc)_2_, 0.1 mM EDTA) and incubated at 4℃ while shaking gently. The concentration of the solution was measured assuming 1 A_260_ = 24 pmol. 15 and 30% sucrose solutions and sucrose gradient was prepared using BioComp GradientMaster^TM^. 300 pmol samples loaded on sucrose gradient taking care that the gradient is undisturbed. The tubes were centrifuged in a Beckman SW41Ti rotor at 21,000 rpm for 16-18 h at 4℃. Fractions were collected and those corresponding to 70S ribosome were dialyzed against buffer C (50 mM HEPES-KOH pH 7.5, 150 mM KOAc, 20 mM Mg(OAc)_2_, 7 mM β-ME, 20 U/mL SuperASE-In RNAse inhibitor).

### CryoEM sample preparation

The dialyzed samples were concentrated using a 100 kDa Amicon® Ultra centrifugal filter, followed by exchange with buffer B (50 mM HEPES-KOH pH 7.5, 150 mM KOAc, 20 mM Mg(OAc)_2_, 7 mM β-ME, 20 U/mL SuperASE-In RNAse inhibitor). The final concentration of the sample was calculated assuming that 1 A_260_ = 24 pmol. Quantifoil R 1.2/1.3, copper, mesh 300 grids with 2 nm amorphous carbon layer on top were glow discharged for 30 s at 15 mA (EMS-100 Glow Discharge System, Electron Microscopy Sciences). 3 µL of a 200-300 nM 70S ribosome sample was applied on the grid and incubated for 30 s at 100% humidity and 4℃. Variable blot times with Whatman #1 filter paper was used to control the ice thickness. Samples were vitrified with a FEI Vitrobot Mark IV (Thermo Fisher).

### CryoEM data collection and processing

The datasets were collected on a Krios 3 electron microscope (200 kV, Thermo Fisher, UCSF cryo-EM core facility) using a nine-show beam image-shift approach with coma compensation in SerialEM and Leginon(20, 21) softwares. The image stacks were collected in non-super resolution mode, binned by a factor of two, motion corrected, and dose-weighted using UCSF motioncor2(22). Dose-weighted micrographs were used to determine contrast transfer function parameters using CTFFIND 4.0 in cryoSPARC(23). Template picker was used to pick particles corresponding to 70S ribosomes(23). The structure 7K00(24) was used to generate a reference map in order to generate 2D classes which were used as input for the template picker. Particles were extracted with a box size of 480 pixels (twice the largest dimension of 7K00). These particles were used for 2D classification. Only classes that clearly contained ice were omitted. Homogeneous refinement was carried out in cryoSPARC using the particles corresponding to the good classes, followed by non-uniform refinement, with both global CTF and defocus refinements turned on(24). 3D classification was carried out twice by dividing the aligned particles from homogeneous refinement into 6 classes with a class similarity of 0.7 to obtain ribosomes in various translational states and filter out noisy particles. The corresponding consensus maps were obtained using cisTEM(25) using identical data processing strategy but using auto refinement with an additional 3D class to remove noisy particles.

### Model building and refinement

The refined and sharpened map obtained after non-uniform refinement in cryoSPARC was used for model building in Coot(26). Restraints for modified nucleotides were generated manually. Model refinement was performed through multiple rounds of manual model building and real space refinement in Phenix(27). We used phenix.real_space_refine to improve the model(27). We used standard Phenix restraints for real space refinement. ModelAngelo was used to model ribosomal proteins(28). The protein residues and nucleotides of the 16S rRNA showed well defined geometrical parameters (Table). The figures were prepared using PyMol Molecular Graphics System Version 3.0.2(29). For *V. cholerae* chimeric ribosomes, we used empty *E. coli* ribosome as a reference (PDB ID 4YBB) due to the current unavailability of native *V. cholerae* ribosome structure.

### Cloning of compensatory mutations and luciferase assays

Compensatory mutants were generated by USER DNA cloning (30), in Mach1F cells, which are Mach1 T1R cells (ThermoFisher Scientific) mated with S2057 F′ to constitutively provide TetR and LacI, as previously described (17). The activity of compensatory mutations was monitored by luciferase assays in the strain S3489, as previously described (17). In brief, single colonies of S3489 strains carrying the o-Ribosome - luciferase genetic circuits were picked to inoculate overnight cultures in Davis Rich Medium (DRM) with with carbenicillin (50□µg□ml^−1^), kanamycin (30□µg□ml^−1^) antibiotic maintenance. Following overnight growth, cultures were diluted 250-fold into fresh DRM supplemented with carbenicillin (50□µg□ml^−1^), kanamycin (30□µg□ml^−1^), and anhydrotetracycline (aTc) (100□ng□ml^−1^) to induce o-Ribosome expression. After 8□h of growth in a 37□°C incubator shaking at 900□rpm., 150□μl of culture was transferred to a 96-well black-wall clear-bottom plate (Costar) and measured for OD600 and luminescence. For kinetic assays, 150□μl of culture was transferred to a 96-well black-wall clear-bottom plate (Costar) 2h after inoculation and measured for OD600 and luminescence every 10 minutes for 8h. In all cases, values were recorded using an Infinite M1000 Pro microplate reader (Tecan) or Spark plate reader (Tecan) running software (SPARKCONTROL version 2.3). Each plasmid combination was assayed in four biological replicates (n = 4).

### Δ-contact map calculation and structure-based visualization

To quantify structural reorganization in mutant ribosomes, we computed changes in intramolecular and intermolecular contacts across rRNA and ribosomal proteins after aligning them in clustalW(31). For each pair of structures (e.g., mutant vs wild-type, or mutant vs chimeric Start), we calculated the difference in contact frequency matrices derived from atomic proximity analysis using a 4□Å heavy-atom cutoff. This yielded separate Δ-contact matrices for RNA-RNA, RNA-protein, and protein-protein interactions. Self-contacts were excluded. Residue-resolved Δ-scores were assigned by summing gains and losses across all interaction partners per residue. Residues with negative Δ-values correspond to net contact loss (destabilization-red), while positive Δ-values indicate net gain (stabilization-blue). The Δ-values for RNA-RNA contact matrices were encoded as B-factors in corresponding PDB files for structure-based visualization in PyMOL using *spectrum b, red_white_blue* command to map contact loss (red) to gain (blue). Residues absent from either structure, or those with local Q-scores below 0.4, were excluded from the analysis to ensure model reliability. To simplify visualization, Δ-contacts were aggregated per nucleotide for RNA and per chain for ribosomal proteins. We focused our structural analysis on regions within 10 Å of the mutation site that are biologically relevant and resolved at a local resolution ≥ 3.0 Å.

## Results

### Structural characterization of evolved ribosomes by CryoEM

To understand how the mutations evolved by oRibo-PACE led to elevated translation kinetics, we recognized that characterization of the corresponding starting ribosomes would be required. In our previous report, 16S rRNA from *Escherichia coli* (starting *E. coli*, EC-ST), *Pseudomonas aeruginosa* (PA-ST; ∼60% activity vs EC-ST), and *Vibrio cholerae* (VC-ST; ∼50% activity vs EC-ST) were used to create chimeras with otherwise wild type host *E. coli* ribosomal proteins and 23S/5S rRNAs (32). These starting ribosomes were evolved to have comparable or higher activities as compared to EC-ST, typified by the 3 prioritized ribosomes EC-S3.5 (∼190% vs EC-ST), PA-S3.3 (∼100% vs EC-ST, ∼160% vs PA-ST), and VC-S4.4 (∼190% vs EC-ST, ∼420% vs VC-ST)(17). In all cases, we structurally characterized the evolved 16S rRNA alleles, where all r-proteins and the 23S/5S rRNAs were derived from the host *E. coli*. Ribosomes were isolated from *E. coli* SQ171(33) strains where all rRNA operons have been replaced by our starting or evolved 16S rRNA chimeras (**Fig. 1A**). PA and VC 16S rRNAs share 85.15% and 90.26% sequence identity to EC-WT 16S rRNA respectively. We note that 9/14 mutations occurred at conserved residues (**Fig. 1B**). Nearly 50% of all mutations occurred in helix 44 (h44), which forms a key element of the ribosomal decoding center(34). Other mutations were found in peripheral regions of the 16S rRNA, many of which have been previously reported to affect translational fidelity(35).

**Figure 1.**
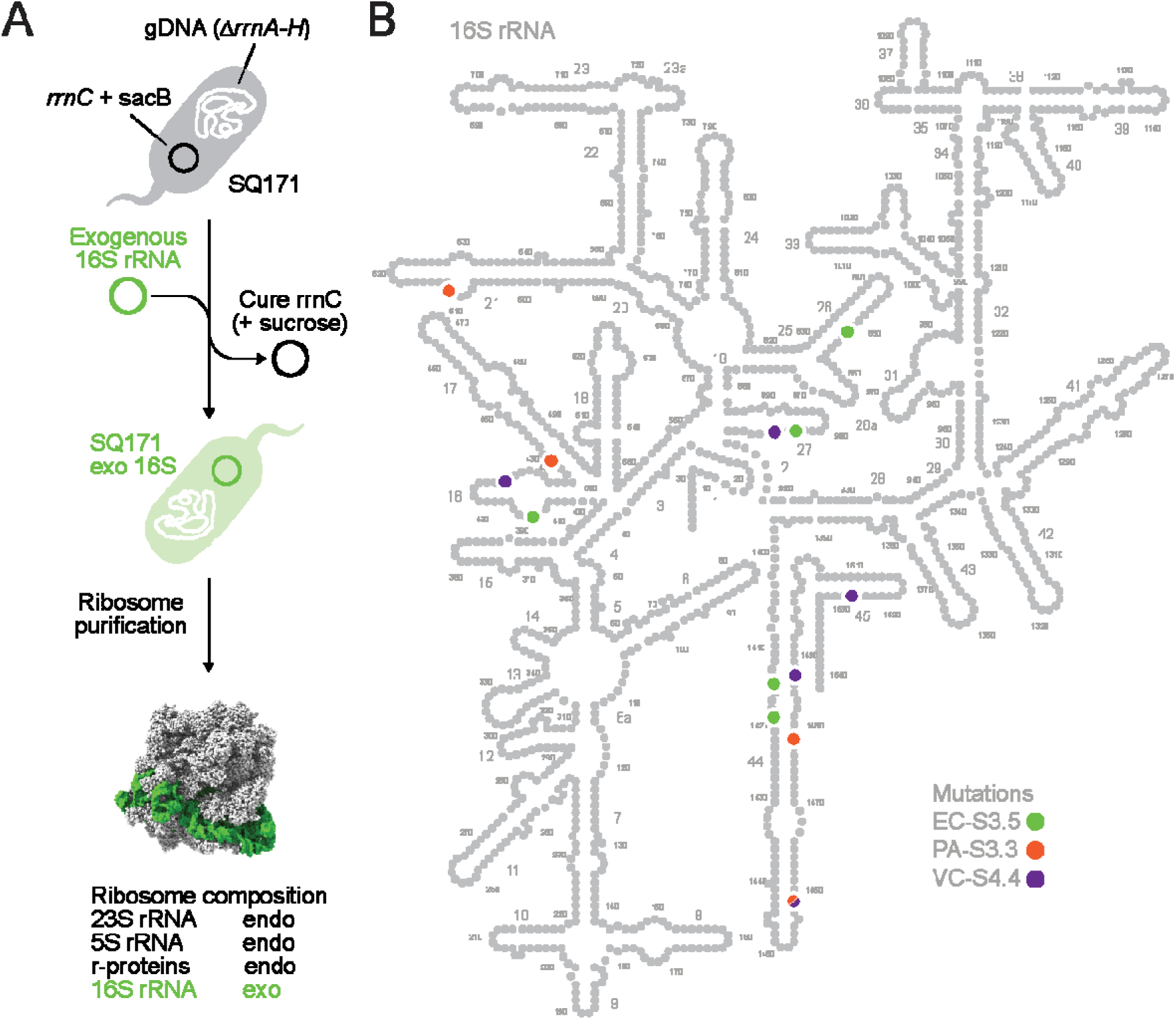
Chimeric ribosomes from o-Ribo-PACE: **A.** A schematic of the *E. coli* SQ171 strain generation indicates the composition of chimeric ribosomes in this study. Exogenous (exo) 16S rRNA is combined with the endogenous (endo) 23S, and 5S rRNA, and r-proteins from an *E. coli* chassis. Throughout the manuscript grey is used to indicate endogenous or wildtype sequence while mutations or impacts of specific mutations are highlighted in color. **B.** The mutations discovered through o-Ribo-PACE, are highlighted on a 16S rRNA map with numbered helices. The 2D 16S rRNA map from *E. coli* (4YBB) highlights mutations as colored circles for *E. coli* (EC-S3.5, green), *P. aeruginosa* (PA-S3.3, orange), and *V. cholerae* (VC-S4.4, purple).

For structural comparisons, different reference models were used depending on the scale of analysis. Local conformational changes at mutated 16S rRNA sites were compared to an established high-resolution empty *E. coli* ribosome structure (PDB 4YBB - 2.10□)(36). Global comparisons of overall 30S architecture and interface packing across chimeric mutants were performed relative to PDB 8G7R (EMD-29821, 2.80□)(37), for which the overall resolution and map quality are most comparable to our reconstructions. PDB 7K00 (1.98□) was included as a high resolution wildtype *E.coli* reference structure only in select cases - to assess the RMSD of EC-S3.5, to compare hydrogen bonds at the uS4-uS5 interface and in the case of VC-S4.4, to compare the conformations of A1492/1493 nucleotides as well as to assess the intersubunit interaction B1c (38) as the region interacting with the 30S subunit in r-protein L31 is unresolved in 4YBB and 8G7R. The wildtype *P. aeruginosa* ribosome structure (PDB 7UNU, EMD-26633, 2.90□)(39) was used as a species-specific reference for PA-derived ribosomes. All ribosomes were purified as previously described(40), and cryo-EM data was processed with cisTEM and cryoSPARC to yield five new ribosome structures with an average resolution of 2.9-3.0Å (**SI Fig. 1, SI Table 1 and 2**). Local resolution analysis revealed a drop off at the periphery, leading us to focus our analysis on regions with resolution better than 3.0Å (**SI Fig. 2**).

### EC-S3.5 mutations are accommodated through local structural adjustments without global rearrangement of 16S rRNA architecture

To globally assess structural changes in our mutant *E. coli*-derived 16S rRNA, we superimposed the EC- S3.5 16S rRNA onto the three highest resolution structures of the *E. coli* ribosome (PDB ID 7K00, 4YBB, 8G7R) (**SI Fig. 3**). The RMSDs ranged from 1.00 to 2.70□ indicating no major global rearrangements beyond the local adaptations described below. The only clear outlier was the crystal structure 4YBB, which shows a slightly more rotated 30S head domain as compared to EC-S3.5, a deviation that may be plausibly influenced by crystallographic packing and crystallization conditions. On the other hand, 7K00 and 8G7R are high-resolution single-particle cryo-EM structures with RMSDs of 0.99 and 2.7□, the latter mostly arising from poorly modeled peripheral regions but with significant overall structural similarity to EC-S3.5. Additionally, we observed increased RNA-protein contacts for EC-S3.5 in comparison with 8G7R (**SI Fig. 4**).

*E. coli* 16S rRNA accumulated mutations in h16, h26, h27, and h44 under oRibo-PACE selections(17). The evolved EC-S3.5 ribosome encodes five 16S rRNA mutations: A412C, G852A, U904C, G1415A, and G1419A. The G1415A mutation occurred during a selection of low mRNA concentration designed to drive evolution towards more efficient translation initiation, while all other point mutants here were found evolving for enhanced elongation rate. Of the five point mutations, G1415A shows the largest impact on the ribosomal structure by realigning the surrounding region in h27, leading to suboptimal stacking interactions in the preceding base pairs U1414-G1486 to optimise hydrogen bonding at the mutation site (**Fig. 2A**). This structural change in h27 displaces uS12, disrupting van der Waals’ contacts within A1413-A1415 and between h44 (U911/U1490) and P90 of uS12 (**Fig. 2A**). uS12 is located at the interface of the 50S and the 30S ribosomal subunits, where it plays a role in rRNA-tRNA-driven movement(41) enforcing translation fidelity(42). We predict that our observed uS12 displacement would affect the volume of the mRNA binding pocket in the decoding center, as P90 and U1490 are critical players in the ribosome’s decoding activity(43, 44).

**Figure 2.**
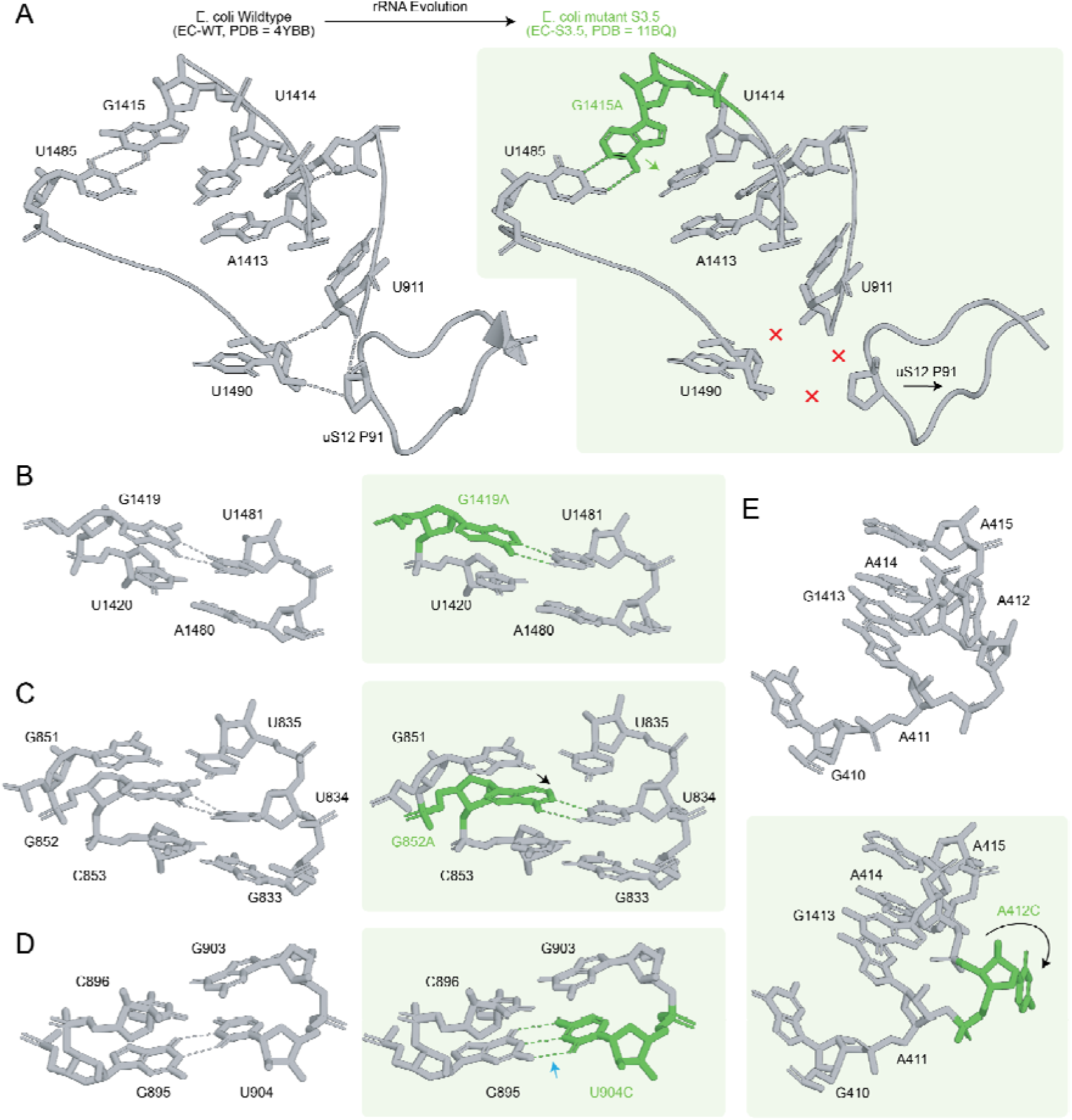
Local structural effects of evolved 16S rRNA mutations in the EC-S3.5 ribosome. Local views of the five evolved 16S rRNA mutation sites in EC-S3.5 compared with empty wild-type *E. coli* ribosome structure EC-WT (PDB ID 4YBB). EC-S3.5 mutation sites are shown in green, and EC-S3.5 panels are shown on a green background. Blue arrows indicate gained contacts and red exes indicate lost contacts in EC-S3.5 when compared with EC-WT. Black arrows indicate the direction of conformational change. **A.** Rearrangement in h44/h27 and the neighboring P90 backbone of uS12 associated with G1415A. **B.** Local rearrangement associated with G1419A in h44. **C.** Local rearrangement associated with G852A in h26. **D.** U904C forms a Watson-Crick G:C base pair in EC-S3.5 instead of the G:U wobble pair observed in EC-WT. **E.** A412C is located in a peripheral, weakly resolved region of the 30S subunit. Dashed lines indicate hydrogen bonds, and arrows indicate the direction of displacement or conformational change.

Realignment of the U1485-A1415 base pair leads to optimization of the stacking interactions between G1415A and U1414. Since the G1415A mutation arose under selective pressure for optimized translation initiation, we sought to investigate whether these structural changes reflect a conformation adapted for this specific step. To test this, we compared our EC-S3.5 structure with the 30S (PDBID 6O7K) and 70S (PDBID 6O9K) initiation complexes. We observed that the backbone conformation of P90 in ribosomal protein uS12 in our structure was more similar to these initiation complexes than to the empty wild-type *E. coli* ribosome (**SI Fig. 5**). This suggests that the G1415A-induced rearrangements may aid initiation by shifting uS12 P90 into an initiation-like open conformation. While these observations provide a plausible structural link to the G1415A phenotype, the relatively low resolution of available initiation complexes (4.20 and 4.00 □ for 6O7K and 6O9K respectively) limits our ability to map the exact differences in hydrogen bonding. Therefore, we caution against drawing definitive mechanistic conclusions regarding the exact step of initiation affected.

The remaining EC-S3.5 mutations resulted in more subtle structural changes. The G1419A and G852A mutations lead to changes from A-U to G-U base pairs in h44 and h26 respectively (**Fig. 2B-C**). These two mutations are likely isoenergetic as both configurations lead to two hydrogen bonds, although optimization of these hydrogen bonds involves displacement of the side chain of the mutated G852A in the direction shown in (**Fig. 2C**). The U904C mutation replaces a G-U by a G-C base pair, increasing the local stabilization of this region, which connects various components of the 30S, such as the decoding center and uS12(38),(45) (**Fig. 2D**). A412 is unpaired in both wild-type and mutant *E. coli* ribosomes (**Fig. 2D**). However, A412C is flipped out in the EC-S3.5 mutant structure when compared to the wildtype *E. coli* ribosome (**Fig. 2E**). Together, these changes indicate that EC-S3.5 accommodates its mutations through minor local base-pair stabilization and repositioning of the side-chains without inducing broader reorganization of the 16S rRNA architecture.

### Accommodation of PA 16S rRNA in EC host leads to additional rRNA-r-protein hydrogen bonds not observed in PA-WT

To understand how chimeric ribosomes with exogenous 16S rRNAs may be structurally altered by *E. coli* ribosomal proteins (r-proteins), we aligned our starting chimeric *P. aeruginosa* (PA-ST) structure to the existing *E. coli* (PDB 8G7R(37)) and *P. aeruginosa* (PDB 7UNU(39)) ribosome structures. We compared the 30S subunit conformation from PA-ST with both structures, by quantifying RNA-RNA, RNA-protein, and protein-protein contacts, which revealed pronounced differences in interface organization (**SI Fig. 6, 7**). Intriguingly, the total number of contacts was higher in PA-ST than both WT structures (**SI Fig. 6, 7**). To minimize confounding effects from low-resolution regions, we next restricted our analysis to well-resolved sites that displayed clear structural deviations. Among these, we quantified significant local differences using thresholds of >10% change in Q-score or >0.5 Å change in local resolution (**SI Tables 3 and 4, SI Fig. 2**). To quantify the significant changes in r-protein-RNA interfaces, we calculated r-protein atoms within 4.0 Å of the 16S rRNA, and found the total number of RNA-protein contacts in PA-ST to be 13.5% and 22.7% higher relative to PDB ID 8G7R(37) and PDB ID 7UNU(39) respectively. This result indicates a modest tightening of rRNA-protein packing. Consistent with this, PA-ST shows a global increase in RNA-protein, RNA-RNA, and protein-protein interactions compared with PA-WT (**SI Fig. 6**). Together, these patterns suggest a subtle constriction of the PA-ST 30S subunit.

To inform our analysis of non-native protein-RNA interactions, we analysed sequence variations between EC and PA r-proteins (**SI Fig. 8**). In general, conservative amino acid substitutions cluster near the buried protein cores, whereas non-conservative substitutions were found primarily at RNA-protein interfaces (**SI Fig. 8A**). This agrees with prior studies showing that RNA-protein interfaces are co-evolutionarily variable(46), and suggests these regions may have played a major role in the structural adaptation of the heterologous 16S rRNAs in our strains. Distinct r-protein conformational differences were observed for a few of *E. coli* r-proteins when accommodating the PA-ST 16S rRNA, most notably bS6, uS8, uS9, uS12 and uS17 (**Fig. 3**). For bS6, the A68Q substitution leads to formation of non-native hydrogen bonding network between R2-R91-Q68 of bS6 and the C733 phosphate in h23 (**Fig. 3A**). For uS8, the S88R substitution introduces an arginine-phosphate salt bridge (R88 NH1 - G627 OP2) while preserving a conserved hydrogen bond (R88 NH2 - G594 OP2) (**Fig. 3B**), likely strengthening the h21-central domain interface(47, 48). We note that bS6 and uS8 stabilize the central domain of 16S rRNA through contacts with helices h21-h23 and coordinate early rRNA folding(49, 50), hinting that these new interactions reinforce early central-domain assembly contacts during accommodation of the heterologous 16S rRNA.

**Figure 3.**
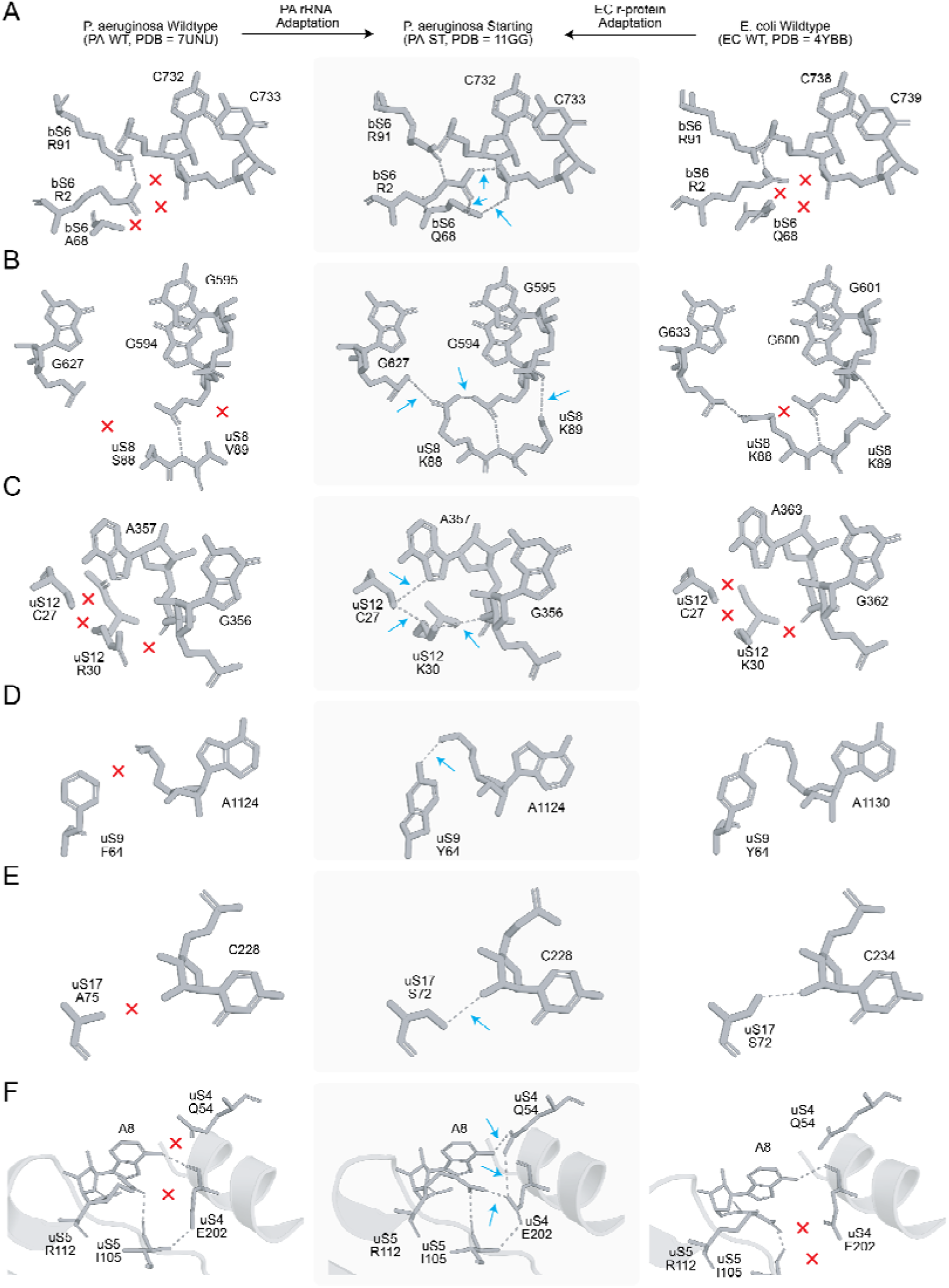
Structural adaptation in *P. aeruginosa* START ribosome: Molecular view of the regions showing the largest conformational changes in the PA chimeric 16S ribosome when compared with PA wildtype (PDB ID 7UNU(39)) and EC wildtype (PDB ID 4YBB(36)) ribosomes. Arrows (blue) mark contacts gained in PA-ST. Exes (red) in the PA-WT and EC-WT panels mark the absence of the corresponding PA-ST contacts in those wild-type structures, highlighting non-native interactions formed in PA-ST. **A.** An A68 (PA) to Q68 (EC) substitution in bS6 leads to formation of non-native contacts at the RNA-bS6 interface. **B.** S88R in uS8 leads to a novel salt bridge between G627 OP2 and R88 NH1. **C.** R30K in uS12 leads to a novel hydrogen bonding network between the NZ atom of K30. **D.** F64Y in uS9 results in a hydrogen bond between uS9 and A1124. **E.** A71S in uS17 results in a hydrogen bond between uS17 and C228. **F.** Molecular view of the non-native interactions with 16S rRNA A8 at the uS4-uS5 interface.

Changes in uS12 occurred at the decoding center, whereas changes in uS9 occurred on the adjacent platform/shoulder region that links the central domain to the decoding center. The uS12 R30K substitution establishes a local hydrogen bonding network involving the amino of K30, the thiol of C27, N7 of A357 and the ribose O3’ of G356 (**Fig. 3C**), reinforcing the RNA pocket adjacent to the decoding center (44). The uS9 F64Y substitution leads to formation of a hydrogen bond between Y64 and A1124, mimicking the interaction present in the *E. coli* wild-type ribosome, where uS9 supports the platform and shoulder regions to connect the central domain to the decoding center (**Fig. 3D**)(51). Finally, uS17 binds the 5’ domain near helices h7-h11 and can stabilize the local rRNA fold at the platform base during assembly of the 30S subunit (51, 52). In the PA-ST ribosome, we observe a non-native hydrogen bond between uS17 S72 and C228 (**Fig. 3E**). Together, these localized protein-RNA interaction changes suggest that *E. coli* r-proteins adapt to the chimeric PA 16S rRNA by increasing local RNA-protein packing and intradomain stabilization, representing early compensatory adjustments that favor accommodation of the heterologous rRNA and integrity of the interface in the PA-ST ribosome.

Apart from these differences, we noted indirect changes in side chain conformations that may have functional implications. We observed three additional hydrogen bonds at the uS4-uS5 interface in the PA-ST ribosome formed by the sidechains of E202 (uS4), R112 (uS5) and I105 (uS5), as well as between A8 of 16S rRNA and Q54 (uS4), which together result in the displacement of the uS5 mainchain by 3.5Å (**Fig. 3F**). To assess the significance of hydrogen bonds between E202, R112 and I105, we examined high resolution models of the *E. coli* (PDB 4YBB - 2.10 □, 5MDZ - 3.10 □, 7K00 - 1.98 □) and *T. thermophilus* (PDB 1VY4 - 2.60 □, 1VY5 - 2.55 □, 1VY6 - 2.90 □, 1VY7 - 2.80 □) ribosomes from the PDB (**SI Fig. 9**). The interprotein hydrogen bonds formed by R112 with E202 and I105 appear only in one of the seven structures (PDB 5MDZ - 3.10 □) that were analyzed and are present in the PA-ST ribosome (**SI Fig. 9**) suggesting that this interaction seen in PA-ST ribosomes is rare. Overall, accommodation of 16S rRNA in the PA-ST ribosome leads to additional rRNA-r-protein hydrogen bonds that do not exist in the PA-WT structure (**Fig. 3**).

### Evolutionary refinement of the PA-S3.3 ribosome through loss of non-native hydrogen bonds

We next investigated the impacts of evolved PA-S3.3 mutations on local 16S rRNA structure. U1472C (PA nucleotide index) transforms a G-U wobble pair into a more stable G-C pair, displacing G1416 and causing a slight rotation to result in optimal hydrogen bonding and stacking interactions (**Fig. 4A, SI Fig. 10-12**). C606U produces the opposite effect, changing a G-C pair to a less stable G-U wobble pair (**Fig. 4B**). In the case of A1453G, an A-U base pair is changed to a G-U pair, causing minimal local change in the side chain due to realignment of U1437 to optimize hydrogen bonding (**Fig. 4C**). In the PA-WT structure, A434 forms a hydrogen bond with U491 at an intrahelical interface region on h16 (**Fig. 4D**), whereas the A434U substitution in PA-S3.3 leads to two supplemental hydrogen bonds with G488 (**Fig. 4D, SI Fig. 10-12**) and the side chain amino group of K121 of uS4 (**Fig. 4D**). We notice PA-ST structure had more contacts than EC WT, but that the evolved PA S3.3 loses a significant number of contacts due to the mutations accumulated during the evolution.

**Figure 4.**
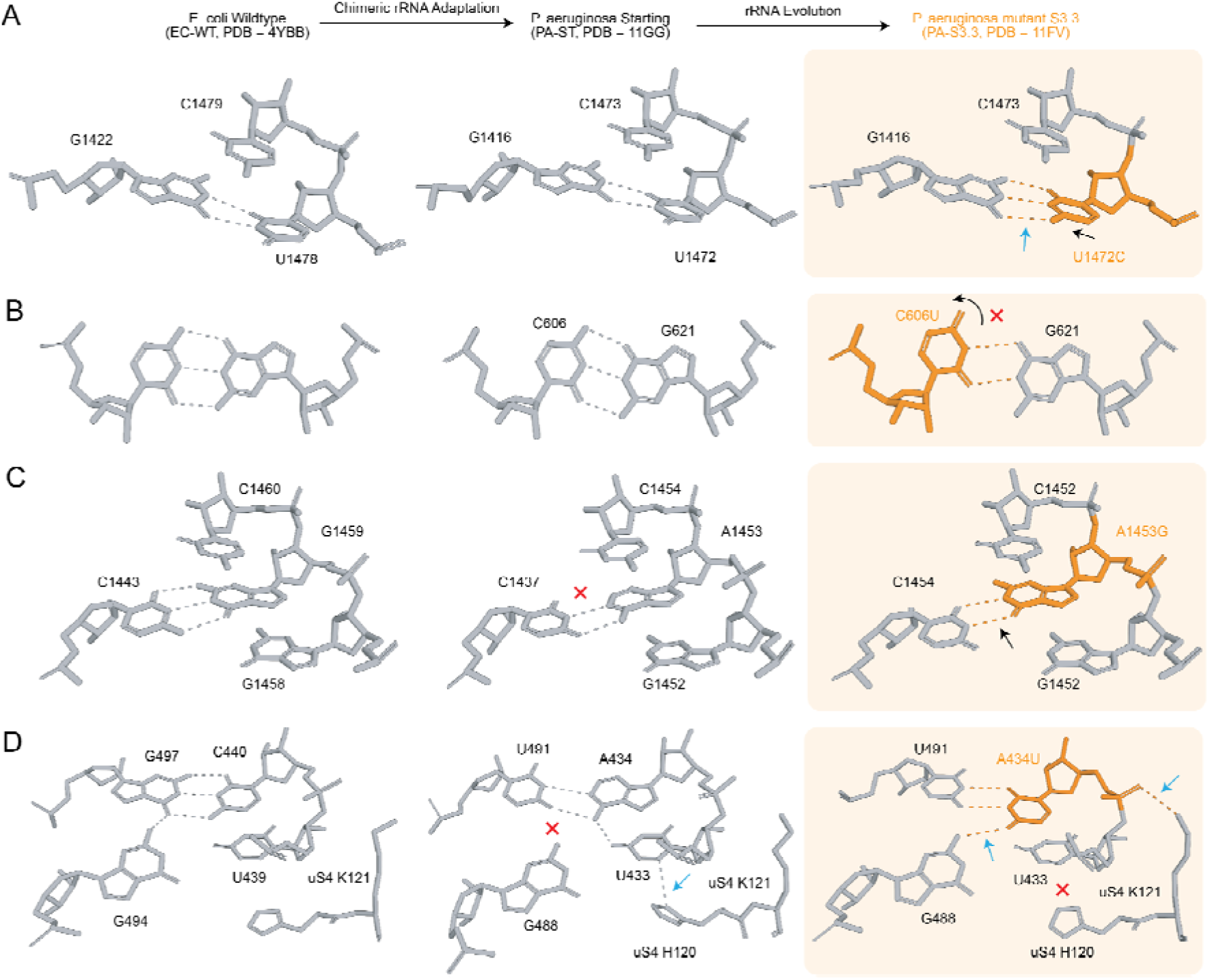
*P. aeruginosa* 16S rRNA mutations impact local stability: Molecular view of the conformational changes associated with mutations in the PA-S3.3 structure. Comparison is made with EC-WT (PDB=4YBB) and PA-WT (PDB=7UNU). All structures are shown (grey) with mutation sites highlighted (orange), and black arrows indicating the direction of conformational change. Blue arrows and red exes indicate gained and lost contacts, respectively, in PA-ST or PA-S3.3 relative to either or both of the reference structures shown in the same row. **A.** U1472C leads to displacement of G1416 and a slight rotation in G1473. **B.** C606U in h21 leads to displacement of the side chain of G622 because of change from a G:C to G:U base pair. **C.** A1453G in h44 causes displacement of U1437 due to a change in the atom that interacts with G1453 from 4’ to 2’-O. **D.** A434U in h16 leads to reorganization of the h16 U434-G488/U491 region and neighboring uS4 interface.

### *V. cholerae* 16S rRNA evolution led to destabilization of h44 and the uS4-uS5 interface

We next analyzed the trajectory that led to one of our most improved orthogonal 16S mutants (compared to the starting point), VC-S4.4. We compared VC-S4.4 with VC-ST (**SI Fig. 13**) and EC-WT (PDB ID 8G7R) (**SI Fig. 14**). No VC-WT comparison was made as no structure exists. Regions of significant structural change (a shift of >1□) were identified by the gain or loss of contacts between RNA-RNA, RNA-protein, and protein-protein, (**SI Fig. 13-14**). We observe an overall reduction in the number of contacts for the VC-S4.4 structure with respect to VC-ST (**SI Fig. 13**), paralleling the adaptation observed along the PA S3.3 trajectory. VC-S4.4 displays a more polarized local resolution profile than VC-ST (**SI Fig. 2**), with higher stability at the peptidyl transferase center and reduced resolution in peripheral regions. In the case of VC-ST, local resolution is comparatively more uniform across the volume. This reflects a reduction in Q-score of VC-S4.4 r-proteins and the 16S rRNA (**SI Fig. 15**). Surprisingly, VC-S4.4 shows better global resolution than VC-ST, suggesting increased conformational heterogeneity (**SI Fig. 2 and 17**). These observations likely indicate destabilization originating from mismatch-induced perturbations at key rRNA-protein interfaces.

Three out of five VC-S4.4 mutations reside on or near h44, which is essential for decoding activity(53). The most prominent local conformational change arises from A1460C (VC nucleotide index) at the apex of h44 (**Fig. 5A**). In VC-ST and EC-WT, A1460 is base-paired to U1444, and A1442 exists in a “flipped out” conformation (**Fig. 5A**). The mutated C1460 pulls A1442 into a “flipped in” conformation, forming a Hoogsteen C1461-A1442 base pair, and introduces a -1 shift in base pairing from nucleotides G1458 to C1461. Poor Q-scores at nucleotides 1442-1446 (**Fig. 5A, SI Fig. 15**) indicate destabilization of this region of h44. C-A base pairs are less stable than the Watson-Crick pair by ∼6.5 kCal/mol and are rare in RNA(54). This destabilization is partially offset by G-U wobble pairs (G1459-U1444 and G1458-U1445). We note that A1460C was not a consensus oRibo-PACE mutation and was found in only 1 of the 32 VC mutants.

**Figure 5.**
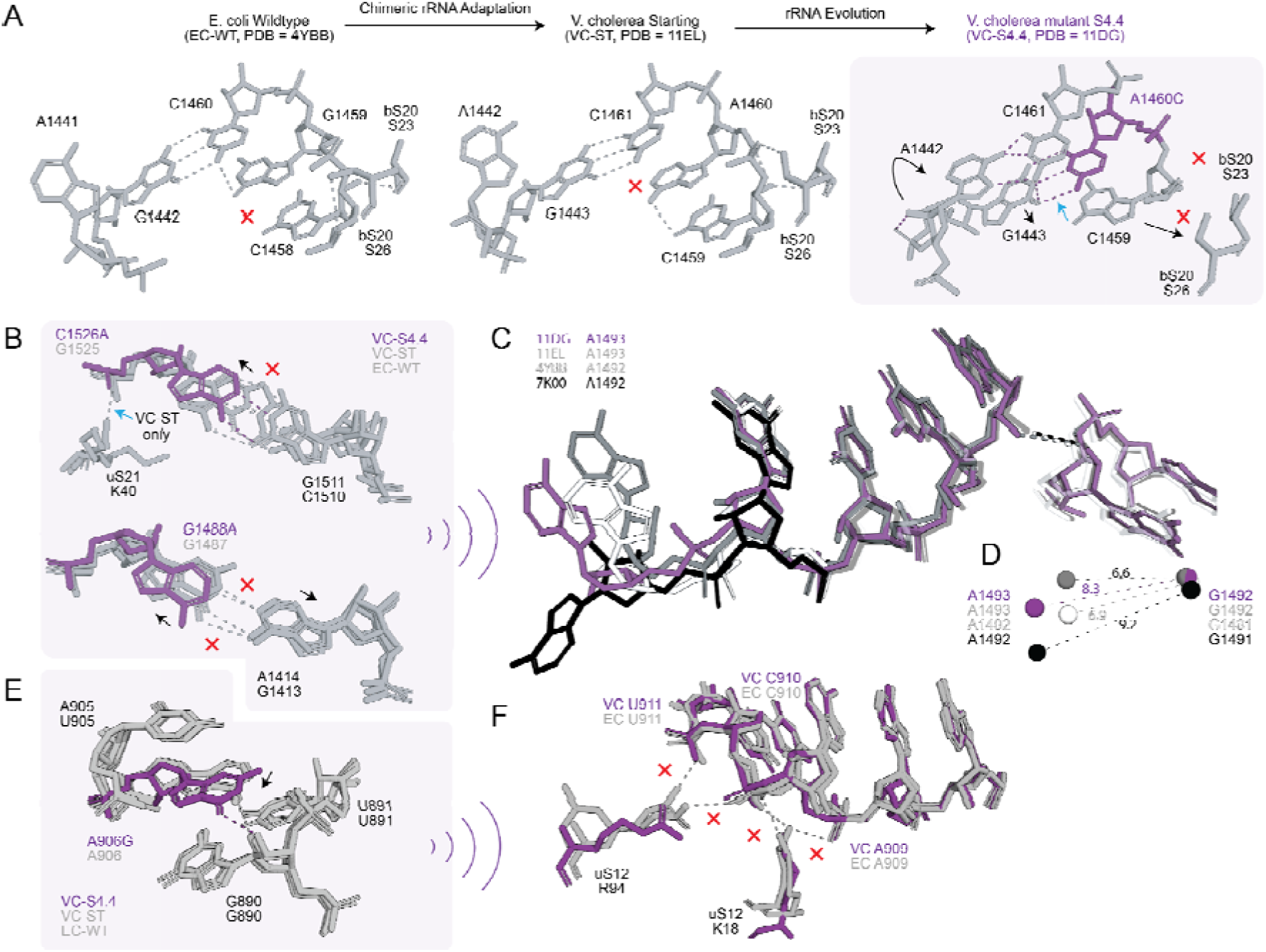
Local h44 and platform mutations in VC-S4.4. The VC ST ribosome is shown in grey, the VC S4.4 ribosome in purple, the empty *E. coli* ribosome (PDB ID 4YBB) in white, and the high-resolution *E. coli* ribosome structure containing an A-site tRNA (PDB ID 7K00) in black. Atoms of the mutated nucleotides are shown in canonical colors. Blue arrows and red exes indicate gained and lost contacts, respectively, in VC-ST and VC-S4.4 in comparison to either or both of the reference structures. Black arrows indicate the direction of displacement or conformational change. **A.** The effect of A1460C (G1459 in *E.coli*) mutation on nucleotides 1442-1447 of h44. The schematic illustrates how the A1460C mutation causes a conformational change in A1442 (A1441 in *E.coli*), which subsequently alters base pairing interactions. **B.** C1526A (G1525 in *E.coli*) mutation leads to the loss of a hydrogen bond with uS21. G1488A mutation leads to the displacement of its own side chain and of A1414 (G1413 in wildtype *E.coli* - PDB ID 4YBB) and **C.** G1488A (G1487) causes local destabilization as a result of the mismatch. A zoomed in view of the conformation of A1493 (A1492 in *E.coli*) in comparison with G1492 (G1491 in *E.coli*). G1488A mutation leads to loss of an intrahelical hydrogen bond between G1489 (G1488 in *E.coli)* and G888 (G888 in *E.coli*) resulting a side chain conformation of A1493 that is intermediate between A-site tRNA bound (PDB ID 7K00) and empty (PDB ID 4YBB) ribosomal states. **D.** Distances between centers-of-mass for the same nucleotides in VC S4.4, VC ST, empty (PDB ID 4YBB) and A-site tRNA bound (PDB ID 7K00) *E.coli* ribosomes. **E**: A906G (A906 in *E.coli*) leads to a slight realignment of its side chain. **F:** Local destabilization due to the mutation A906G leads to the loss of hydrogen bonds between uS12 (Arginine (R) 94 and Lysine (K) 18) and C910 (C910 in *E.coli*).

We also observed a subtle change at the 30S-50S interface linked to the altered conformation of A1442. In VC-ST and EC-WT (PDB ID 4YBB), A1442 forms a CH-π interaction with L114 of 50S protein L19, which is disrupted upon A1442 flipping in VC-S4.4 (**SI Fig. 16A**). We additionally detected disruption of two intersubunit contacts outside the mutated regions. First, the B1c bridge, involving E65 and H69 of uS19, R79 of uS13, and R56 of 50S protein L31 (38), is intact in VC-ST and EC-WT (PDB ID 7K00), but disrupted in VC-S4.4 (**SI Fig. 16B**). Although density for L31 R56 is weak in VC-S4.4, the remaining side chains could be modeled and support loss of the associated hydrogen-bonding network. PDB ID 7K00 was used as the EC-WT reference for this comparison because the R56-containing region of L31 is not modeled in either 4YBB or 8G7R. Second, the B4 bridge, involving Q40 of uS15 and A715/A716 of 23S rRNA (38), is also disrupted in VC-S4.4 but retained in both EC-WT (PDB ID 4YBB) and VC-ST (**SI Fig. 16C**).

Additional mutations, C1526A and G1488A, led to realignment of bases that optimizes hydrogen bonding for both the mutated nucleotides (**Fig. 5B**). The C1526A mutation changes base pairing from a stable G-C to a relatively unstable G-A pair, which causes displacement in both side chain and phosphate backbone. The displacement disrupts a hydrogen bond between K40 in uS21 r-protein and the nucleotide 1526 (**Fig. 5B**), which would add to the destabilization of the body domain of 16S rRNA in the VC-S4.4 ribosome. The G1488A mutation, on the other hand, causes a change from a relatively unstable G-A base pair to highly unstable A-A clash resulting from hydrogens of A1488 and A1414 (**Fig. 5B**).

We observe that the backbone of nucleotides A1492 and A1493 (A1493 and A1494 in VC) is displaced. Since A1492 and A1493 are important for tRNA binding at the A-site, we compared our structure with the high-resolution A-site tRNA bound structure (PDBID 7K00) to compare the conformations adopted by these nucleotides (**Fig. 5C**). The displacement of the A1488 side chain positions A1493 (A1492 in *E. coli*) in a partially open conformation intermediate between the empty EC-WT/VC-ST and A-site tRNA-bound EC-WT (7K00) structures (**Fig. 5C**). This is reflected in the distance between the centers of mass of A1493 (A1492) and G1492 (G1491), which is 8.3 Å in VC S4.4. This value is between 9.2Å for the open, tRNA-bound state (PDB ID 7K00) and 6.6 - 6.9 Å for the closed state (VC-ST and empty EC-WT, PDB ID 4YBB) (**Fig. 5D**). A906G mutation leads to a minimal local displacement of its side chain (**Fig. 5E**), leading to the disruption of hydrogen bonds between K18 and R94 of uS12 with nucleotides 909-911 which remain intact in the EC-WT structure (**Fig. 5F**). The hydrogen bond between the 2’-OH of G890 and the mutated G906 on the other hand, is retained in all three structures (VC-S4.4, EC-WT, and VC-ST) (**Fig. 5E**). K18 lies within the N-terminal extension of uS12, which is required for proper 30S assembly and maintenance of local 16S rRNA architecture in the platform region (55), whereas R94 resides in the conserved C-terminal uS12 loop that harbors classic *rpsL* mutations known to modulate translational output by perturbing the decoding-site environment (44, 56, 57). Loss of K18/R94 contacts to nucleotides 909-911 therefore may alter the local uS12-rRNA interface near the decoding center. Together with mutations G1488A, C1526A, and A1460C, which introduce both local and long-range changes in h44, and the more localized effects of A906G on uS12-rRNA contacts at the base of the helix, these structural changes correlate with the increased ribosome activity of the evolved VC S4.4 ribosome.

Due to the proximity of U426C (h16) and uS4, we believe a large perturbation observed at the uS4-uS5 interface may be attributed to the loss of the K33 uS4 - U426 interaction (**Fig. 6A**). The U426C mutation introduces an additional hydrogen bond at the C426-G417 pair relative to the native U426-G417 interaction, resulting in local stabilization of the base-pair (**Fig. 6B**). However, realignment of the nucleotide side chain due to the new G-C base pair shifts the phosphate backbone leading to a breakage of hydrogen bonds formed by K33 and Q36 of uS4 with U426 (**Fig. 6B**). This perturbation manifests as displacement of the uS4 main chain and loss of multiple hydrogen bonds at the uS4-uS5 interface.

**Figure 6.**
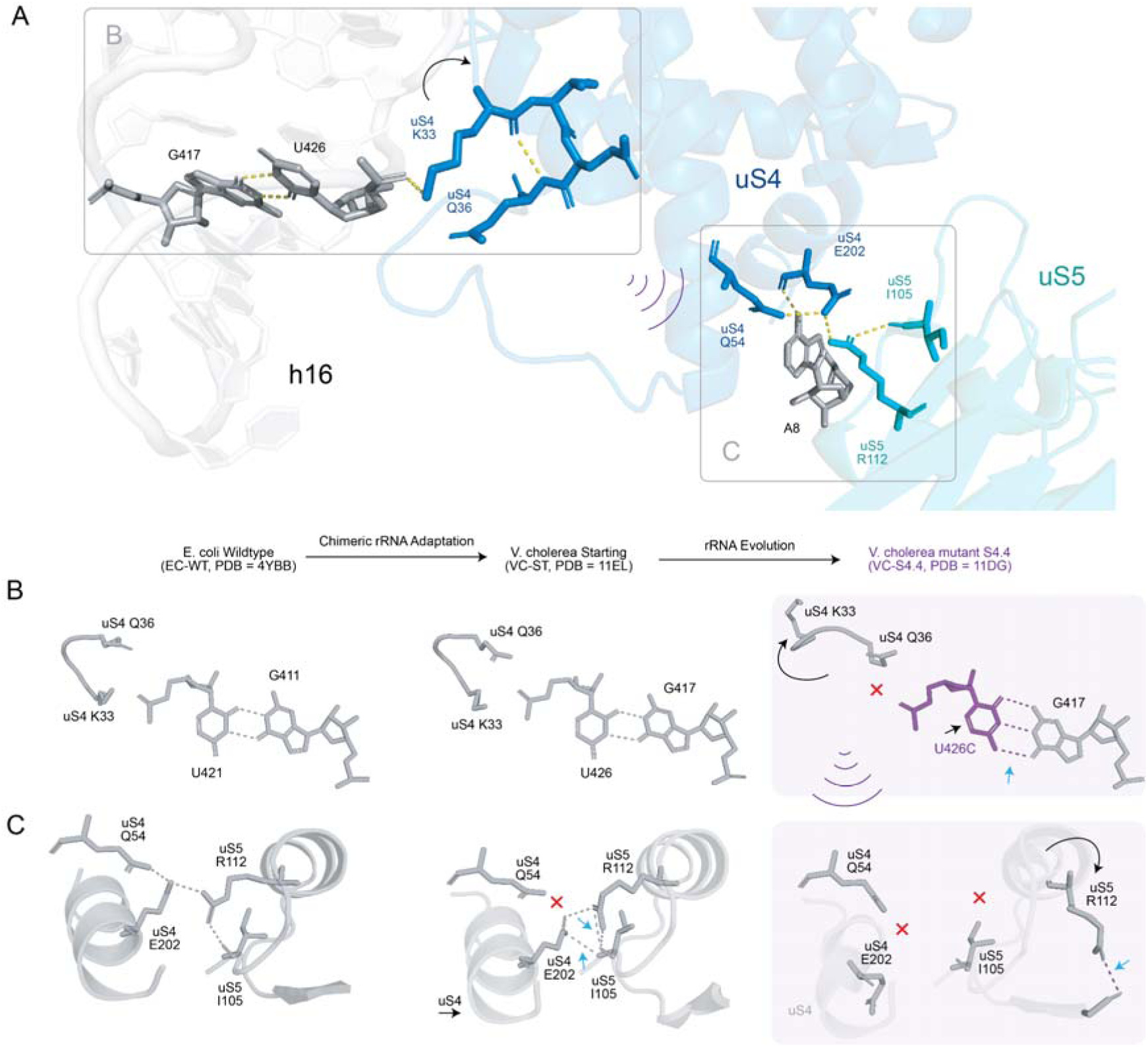
Impacts of VC-S4.4 U426C mutation on the uS4-uS5 interface. All structures are shown (grey) with mutation sites highlighted (purple). Blue arrows and red exes indicate gained and lost contacts, respectively, in VC-ST and VC-S4.4 in comparison to either or both of the reference structures. Black arrows indicate the direction of displacement or conformational change. **A.** Molecular view showing the proximity of 16S rRNA h16 and A8 to the uS4-uS5 interface. U426 and G417 are shown as sticks (grey) on h16. Relevant residues in uS4 (blue) and uS5 (teal) are shown as sticks, while h-bonds are shown as dashed lines (yellow). **B.** Potential allosteric effect of the U426C mutation resulting from the re-matched C426:G417 base pair leading to conformational change in the backbone of uS4 (Lysine (K) 33 and Glutamine (Q) 36), **C.** U426C mutation causes the destabilization of the uS4-uS5 interface (color spheres). Here, the uS4-uS5 interface in VC chimeric S4.4 (purple) is superposed with the VC-ST (Grey, aligned to uS4) ribosome.

Impacted inter-protein contacts include those between uS4 E202 and uS5 I105, uS4 E202 and uS5 R112, and between A8 and uS4 Q54 (**Fig. 6C**). Interestingly, both VC-ST and PA-ST chimeric ribosome structures adopt a similar uS4-uS5 stabilization mechanism. VC-S4.4 shows a greater displacement and rotation of the 30S subunit compared to VC-ST (11° vs 7°; **SI Fig. 17**). Additionally, VC-S4.4 dataset was classified into multiple conformational states (**SI Table 2**). Our results showing the breakage of these specific hydrogen bonds at the uS4-uS5 interface in our structures is consistent with the disruption of this interface during the open-to-closed transition of the ribosome(58).

We note that class occupancies in (**SI Table 2**) are influenced by several factors such as image-processing and experimental parameters such as ice thickness, sample heterogeneity, particle orientation distributions, optical aberrations, local resolution. As a result, these class occupancies are not interpreted quantitatively but the presence of several discrete classes qualitatively supports enhanced conformational sampling in VC-S4.4. The conformational destabilization of the uS4-uS5 neck region due to U426C is consistent with elevated translational output for VC-S4.4 as this interface mediates head-body motions required for mRNA accommodation (59), (60).

### Structure-guided ribosomal RNA engineering can restore translation levels

A notable feature of both PA-S3.3 and VC-S4.4 was local changes in contacts within h44. These differences are correlated with local destabilization in this region and we hypothesized that they contribute to the increased activity observed in the evolved PA and VC mutant ribosomes. As a forward test of this hypothesis, we designed compensatory mutations that restore canonical base pairing at the mismatched positions A1453G in PA-S3.3 and A1460C in VC-S4.4 (**Fig. 7**). In PA-S3.3, the A1453G substitution occurs in the central portion of h44, where it converts a native A-U Watson-Crick pair into a G-U wobble pair (**Fig. 7A**). Based on known thermodynamic comparisons, this substitution is expected to be only mildly destabilizing relative to the canonical A-U interaction(61, 62). To restore Watson-Crick pairing at this position, we introduced the U1437C compensatory mutation, which re-establishes a G-C base pair. In VC-S4.4, the A1460C substitution occurs at the apex of h44, where it disrupts the native A1460-U1444 Watson-Crick pair and introduces a local base-pairing rearrangement (**Fig. 7B**). Like the A1453G change in PA-S3.3, this mismatch is consistent with local destabilization of this segment of h44. To restore Watson-Crick pairing at this position, we introduced the U1444G compensatory mutation, which restores C1460 - G1444 pairing and prevents downstream disorder. Next, we tested the functional effect on ribosome activity. Both compensatory mutants showed lower activity than their evolved PA and VC counterparts, indicating that restoring Watson-Crick pairing and thereby locally stabilizing h44 reduces activity. Together, these results support the interpretation that a higher degree of local destabilization in h44 is associated with increased translational activity in the chimeric PA-S3.3 and VC-S4.4 ribosomes.

**Figure 7.**
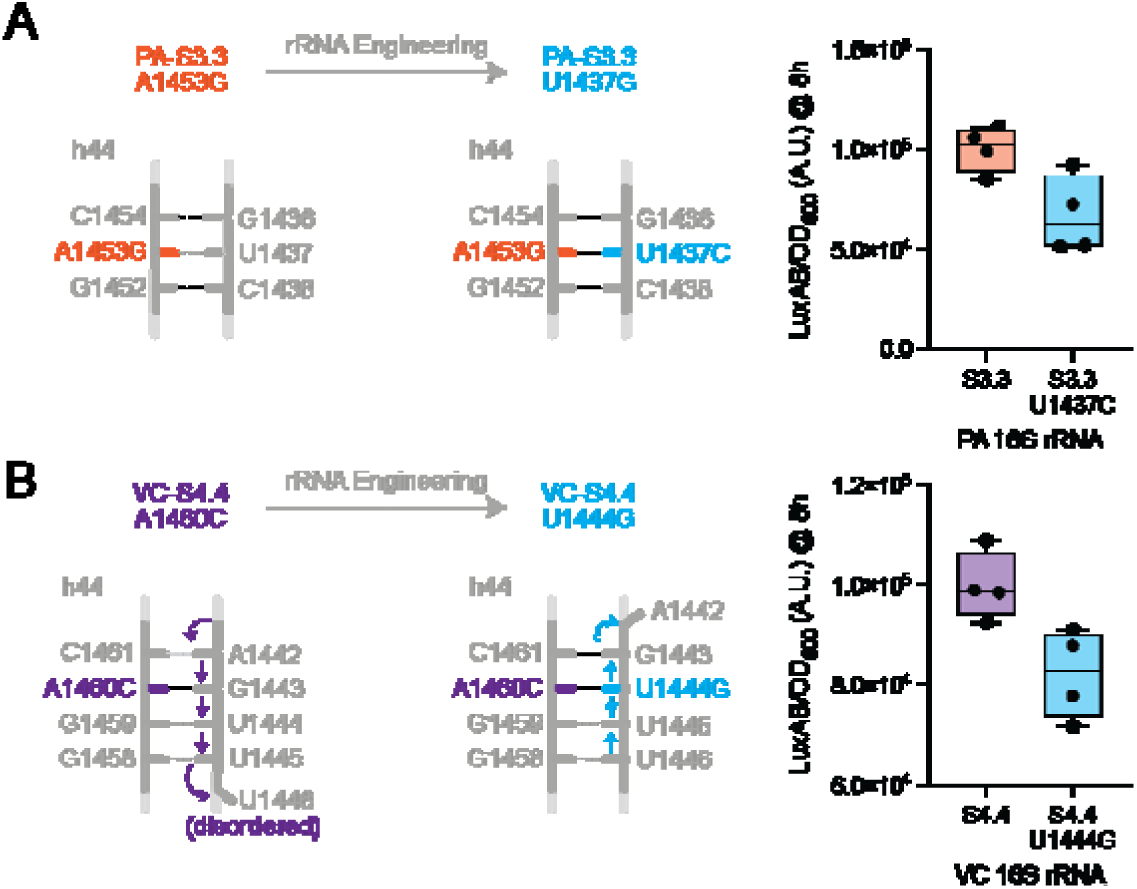
Model for structural adaptation and increased activity in chimeric ribosomes. Compensatory mutations were introduced to PA-S3.3 and VC-S4.4 16S rRNA to stabilize mismatch bases discovered through rRNA evolution. **A.** PAIJ-S3.3, the A1453G substitution converts an A-U Watson Crick pair to a G-U wobble pair. A U1437C compensatory mutation was introduced (G-C base pair), to restore Watson Crick pairing. The translational activity of PA-S3.3 (orange) and engineered variant (blue) were monitored by luciferase assay, (n = 4). **B.** The A1460C mutation in VC-S4.4 causes a base pairing rearrangement in h44. Here we designed a U1444G compensatory mutation to re-match the C1460 to G1444 and avoid the subsequent disordering of base pairs. Translational activity of VC-S4.4 (purple) and the compensatory engineered variant (blue) were monitored by luciferase assay, (n = 4). Data is plotted as a box and whiskers plot where the whiskers represent the min and max of all data points.

## Discussion

In this work, we investigated the structural adaptation of three 16S rRNA species following evolution with oRibo-PACE. In these experiments, orthogonal 16S rRNAs were evolved for increased activity while operating with *E. coli* 23S/5S rRNAs and ribosomal proteins. The resulting chimeric ribosomes reveal distinct patterns by which heterologous rRNA is accommodated in starting and evolved variants. We hypothesized that the introduction of exogenous 16S species could present with sub-optimal translational activity due to improper RNA folding or disruption of RNA-RNA and RNA-protein interfaces. Our observations suggest that chimeric ribosomes (PA-ST and VC-ST) form additional RNA-protein contacts due to subtle structural changes in 16S o-rRNA, with differences in PA and VC r-protein surfaces relative to EC enabling these additional interactions. Many of these contacts are known to be essential for proper assembly of the 30S subunit (55, 63, 64). As a result, these additional contacts may be constricting and explain the loss in translational activity of these ribosomes.

However, those constricting interactions can be compensated for with subsequent mutations enriched by selection. We find our evolved ribosomes exhibit extensive structural adaptations, where local base substitutions lead to large RNA rearrangement or disruption of RNA-protein interfaces. Indeed, one of our most active orthogonal ribosome compositions across various phenotypic assays, VC-S4.4, found an evolutionary trajectory to break non-native contacts formed in the chimeric VC-ST structure. VC-S4.4 acquires the mutation A1460C which causes a local perturbation at the tip of h44. Here, A1460C forms a non-native C-G pair with G1443 causing the formation of a C-A Hoogsteen base pair and leads to a rearrangement of nucleotides 1442-1446. Beyond this, we find a subtle change in local base pairing between U426C and G417 in h16 disrupts a native contact with uS4 K33. This minor base-pairing optimization appears to have a large cascading effect on the uS4-uS5 interface. Together, these changes may broaden the conformational ensemble sampled by the 30S subunit and correlate with the higher translational activity we observe in phenotypic assays with this ribosome. When considered alongside phenotypic assays, these observations suggest that mismatched interfaces and selective local loss of contacts can have divergent functional consequences.

Our results suggest that the interfaces in the ribosome are not optimized for any single conformation due to the multiple conformational rearrangements that are needed for the translational cycle. For example, the uS4-uS5 interface contributes to mRNA entry and positioning at the decoding center(65). In PA-ST, the presence of three additional hydrogen bonds stabilizes this interface relative to EC-WT. This increased stabilization may restrict local flexibility at the neck region and is therefore consistent with reduced decoding efficiency in PA-ST. The reduced translational speed of the designed mutations further support this hypothesis. Although our structures reveal clear patterns of local accommodation and disorder, several limitations temper mechanistic interpretation. Each evolved ribosome carries multiple dispersed mutations, and we did not obtain structures of individual point variants, making it difficult to disentangle additive contributions from those that are epistatic. Many substitutions generate only subtle local rearrangements while downstream regions shift more substantially, meaning some inferred allosteric effects cannot be assigned with precision. Likewise, the increased conformational heterogeneity observed in VC-S4.4 arises from qualitative 3D classifications, rather than quantitative dynamic measurements, limiting direct conclusions about altered motions. Finally, we acknowledge the limitations associated with our phenotypic characterization, namely the use of reporter assays to infer ribosome activity in most cases, where only a subset of 16S rRNA mutants have undergone more in-depth evaluation for global protein synthesis (AHA incorporation rate), and assessment of in vitro translational fidelity. Further insights gained through the assessment of in vitro translation kinetics or mutant 16S biogenesis may help to improve our understanding going forward.

Despite these constraints, our findings highlight a set of peripheral, structurally tolerant regions of 16S rRNA as promising “designable zones” where ribosomal behavior can be tuned without perturbing core decoding architecture. Integrating such structural insight with in vivo selections like oRibo-PACE will enable further engineering of orthogonal ribosomes with tailored dynamics, enhanced orthogonal translation activity, or decoding preferences. More broadly, coupling targeted mutagenesis with high-resolution structures and genotype-phenotype maps may enable predictive design of ribosomes with user-defined properties.

## Data and code availability

Cryo-EM maps and coordinates generated in this study have been deposited in the Protein Data Bank (PDB) and the Electron Microscopy Data Bank (EMDB). The following structures were deposited as part of this study: EC-S3.5 (PDB: 11BQ, EMDB: EMD-75605), PA-ST (PDB: 11GG, EMDB: EMD-75676), PA-S3.3 (PDB: 11FV, EMDB: EMD-75668), VC-ST: (PDB: 11EL, EMDB: EMD-75650), VC-S4.4 (PDB: 11DG, EMDB: EMD-75634). Scripts used to compute RNA-RNA, RNA-protein and protein-protein contact difference matrices and to generate the corresponding heat maps are available at: https://github.com/rtushar76/Difference-contact-map

## Supplementary data

Supplementary Data are available at *NAR* Online

## Acknowledgements

We are grateful for the help of the UCSF EM core (Dr. David Bulkley and Glenn Gilbert) and the staff of the New York Structural Biology Center for their assistance in data collection and analysis. We thank Justin Sim and Rishav Mitra for their helpful discussions and comments.

## Author contributions

T.R. & A.C. designed the study and contributed to experimental work. T.R. performed sample preparation, cryo-EM data collection and analysis. T.R. and A.C. wrote the manuscript with input from all authors. A.H.B and J.S.F conceived and supervised the project. All authors reviewed, commented on and approved the final paper.

