## Supplemental Information for "Structural adaptations for enhanced translation kinetics in evolved ribosomes"

### Supplementary Information

#### **Supplementary Table 1.**

Cryo-EM data collection, processing, and model refinement statistics (provided separately as a spreadsheet)

#### **Supplementary Table 2.**

Particle and resolution statistics for the respective volumes obtained using the data processing strategy in this study (SI Fig. 1B).

|  | **EC-S3.5** | **PA-ST** | **PA-S3.3** | **VC-ST** | **VC-S4.4** |
| --- | --- | --- | --- | --- | --- |
|  | Number of particles  (resolution after homogeneous refinement) | | | | |
| **Homogeneous refinement**  **(all particles)** | 563168  (2.8 Å) | 570940  (3.0 Å) | 997398  (2.6 Å) | 1210610  (3.0 Å) | 746202  (2.7 Å) |
| **Class I** | 86066  (3.1 Å) | 125041  (3.8 Å) | 75503  (3.0 Å) | 323557  (3.5 Å) | 249657  (2.9 Å) |
| **Class II** | 83903  (3.5 Å) | 151017  (3.5 Å) | 169289  (3.1 Å) | 111810  (3.4 Å) | 138707  (3.3 Å) |
| **Class III** | 224193  (3.2 Å) | 8169  (6.3 Å) | 107892  (3.2 Å) | 269266  (3.2 Å) | 53500  (3.3 Å) |
| **Class IV** | 64169  (3.1 Å) | 49049  (5.9 Å) | 77069  (3.1 Å) | 89201  (3.5 Å) | 116432  (3.1 Å) |
| **Class V** | 95757  (3.3 Å) | 202321  (5.9 Å) | 423509  (2.9 Å) | 230089  (3.2 Å) | 70530  (3.1 Å) |
| **Class VI** | 9080  (4.7 Å) | 35343  (3.8 Å) | 145136  (3.0 Å) | 186685  (3.3 Å) | 117377  (3.2 Å) |

#### **Supplementary Table 3.**

**Q-scores for ribosomal proteins from all datasets.**

| **r-protein** | **EC-S3.5** | **PA-ST** | **PA-S3.3** | **VC-ST** | **VC-S4.4** |
| --- | --- | --- | --- | --- | --- |
| **uS2** | 0.62 | 0.44 | 0.62 | 0.67 | 0.57 |
| **uS3** | 0.67 | 0.71 | 0.70 | 0.69 | 0.63 |
| **uS4** | 0.68 | 0.67 | 0.70 | 0.70 | 0.64 |
| **uS5** | 0.72 | 0.73 | 0.75 | 0.73 | 0.70 |
| **bS6** | 0.67 | 0.68 | 0.65 | 0.70 | 0.36 |
| **uS7** | 0.56 | 0.68 | 0.65 | 0.64 | 0.53 |
| **uS8** | 0.72 | 0.74 | 0.76 | 0.73 | 0.71 |
| **uS9** | 0.61 | 0.69 | 0.63 | 0.68 | 0.60 |
| **uS10** | 0.58 | 0.60 | 0.69 | 0.62 | 0.55 |
| **uS11** | 0.69 | 0.71 | 0.70 | 0.71 | 0.66 |
| **uS12** | 0.73 | 0.68 | 0.74 | 0.73 | 0.70 |
| **uS13** | 0.58 | 0.70 | 0.69 | 0.69 | 0.51 |
| **uS14** | 0.64 | 0.67 | 0.60 | 0.68 | 0.64 |
| **uS15** | 0.71 | 0.70 | 0.74 | 0.72 | 0.58 |
| **bS16** | 0.63 | 0.65 | 0.72 | 0.54 | 0.66 |
| **uS17** | 0.58 | 0.69 | 0.67 | 0.68 | 0.67 |
| **uS18** | 0.69 | 0.72 | 0.73 | 0.70 | 0.52 |
| **uS19** | 0.59 | 0.58 | 0.62 | 0.66 | 0.56 |
| **uS20** | 0.69 | 0.68 | 0.73 | 0.65 | 0.67 |
| **uS21** | 0.59 | 0.56 | 0.54 | 0.62 | 0.44 |
| **16S rRNA** | 0.68 | 0.70 | 0.74 | 0.70 | 0.65 |

#### **Supplementary Table 4.**

**Net gains and losses of protein-protein, RNA-protein, and RNA-RNA contacts in ribosome structures based on difference contact plot analysis:**
The values represent the signed counts of atomic contacts computed by subtracting the contact counts of the reference structure from those of the mutant or alternative structure. Positive values indicate a net gain of contacts in the comparison, while negative values indicate a net loss. Protein-protein contacts are reported separately for inter-protein interfaces and the combined total of intra and inter-protein contacts. RNA-protein and RNA-RNA contacts were calculated by applying residue-level Q-score filtering prior to contact counting to ensure only well-resolved regions contributed to the analysis.

| **Comparison** | **Inter-protein contacts (off-diagonal)** | **Intra-protein contacts (on-diagonal)** | **RNA-protein contacts** | **RNA-RNA contacts** |
| --- | --- | --- | --- | --- |
| **EC-S3.5 - EC-WT** | 70 | 1274 | 700 | 21716 |
| **PA-S3.3 - EC-WT** | 14 | 72 | -506 | -4570 |
| **PA-S3.3 - PA-ST** | -56 | -1026 | 100 | -23812 |
| **PA-S3.3 - PA-WT** | -12 | -482 | -801 | -10462 |
| **PA-ST - EC-WT** | 70 | 1116 | 1623 | 19218 |
| **PA-ST - PA-WT** | 41 | 618 | 928 | 13780 |
| **VC-S4.4 - VC-ST** | -69 | -1778 | -1685 | -24130 |
| **VC-S4.4 - EC-WT** | 0 | -410 | -35 | -6650 |
| **VC-ST - EC-WT** | 86 | 1514 | 1742 | 17986 |

#### SI Figure 1

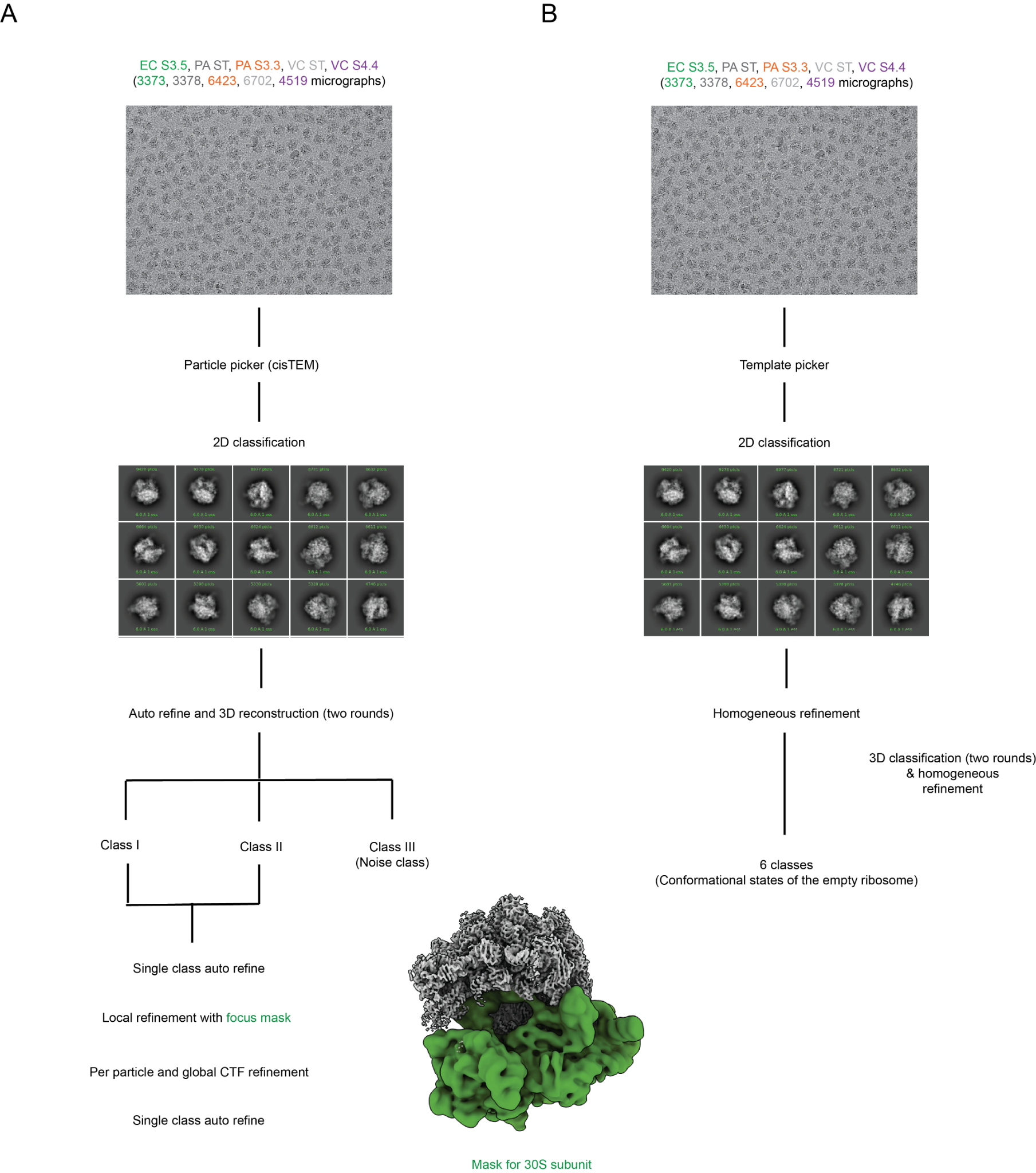

**Supplementary Figure 1. Data processing workflow:**

**A.** The workflow to generate a consensus volume for model building and refinement.

**B.** The workflow used for 3D classification.

Raw data were processed using cryoSPARC and cisTEM (23),(25). cisTEM was used to generate consensus volumes, and cryoSPARC for 3D classification. The volumes obtained in cisTEM were then used for Real Space Refinement in Phenix(66), followed by manual model building in COOT (67).

#### SI Figure 2

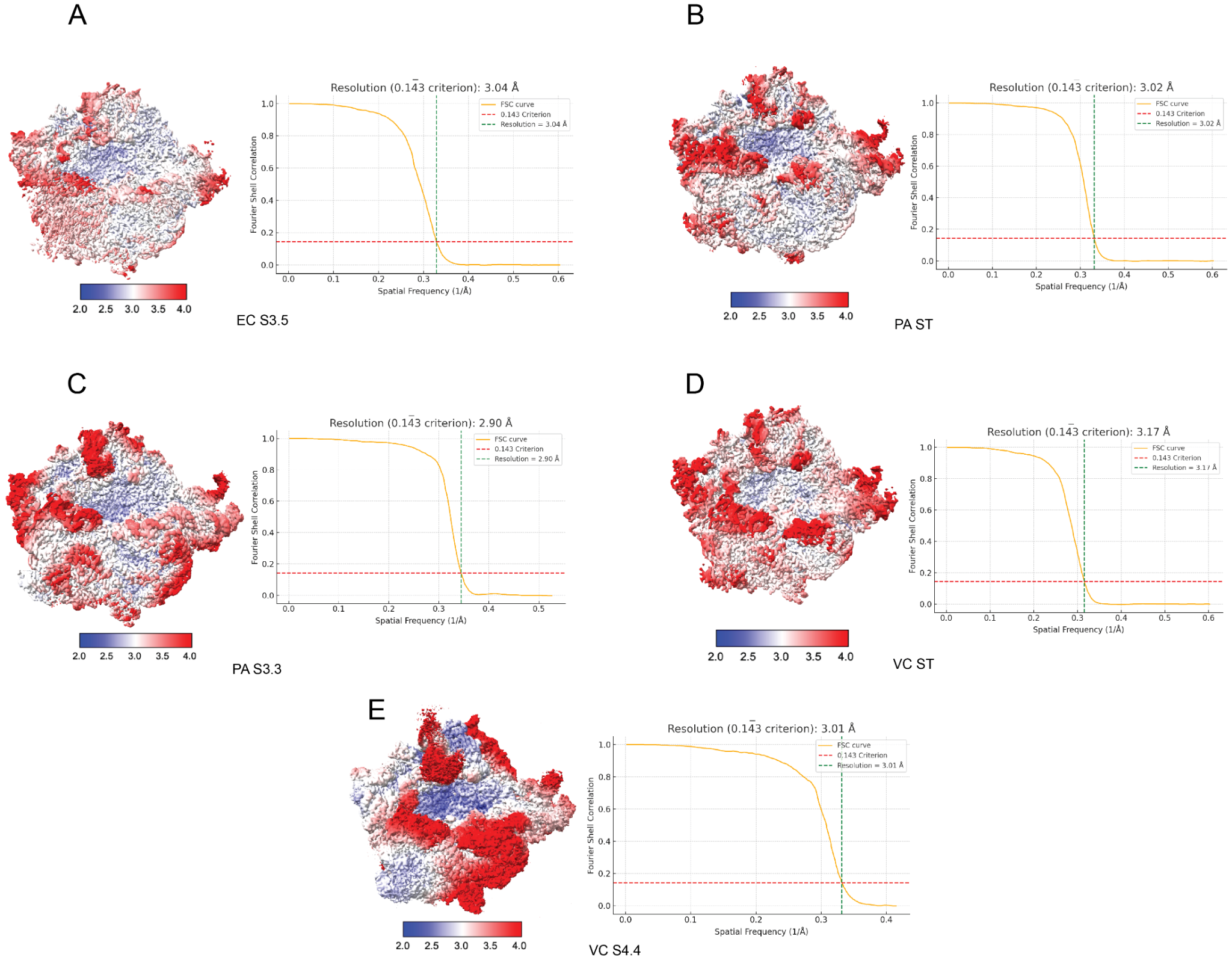

**Supplementary Figure 2. Local resolution calculation:**

Fourier shell correlation curve for maps obtained using homogeneous refinement for

**A.** EC-S3.5,

**B.** PA-ST,

**C.** PA-S3.3,

**D.** VC-ST, and

**E.** VC-S4.4 datasets.

The average resolutions for these maps were found to be between 2.90 and 3.17Å (unmasked) as indicated by the horizontal red and the vertical green lines**.** The 70S ribosome map is colored by the local resolution for the consensus map obtained from homogeneous refinement in cisTEM.

#### SI Figure 3

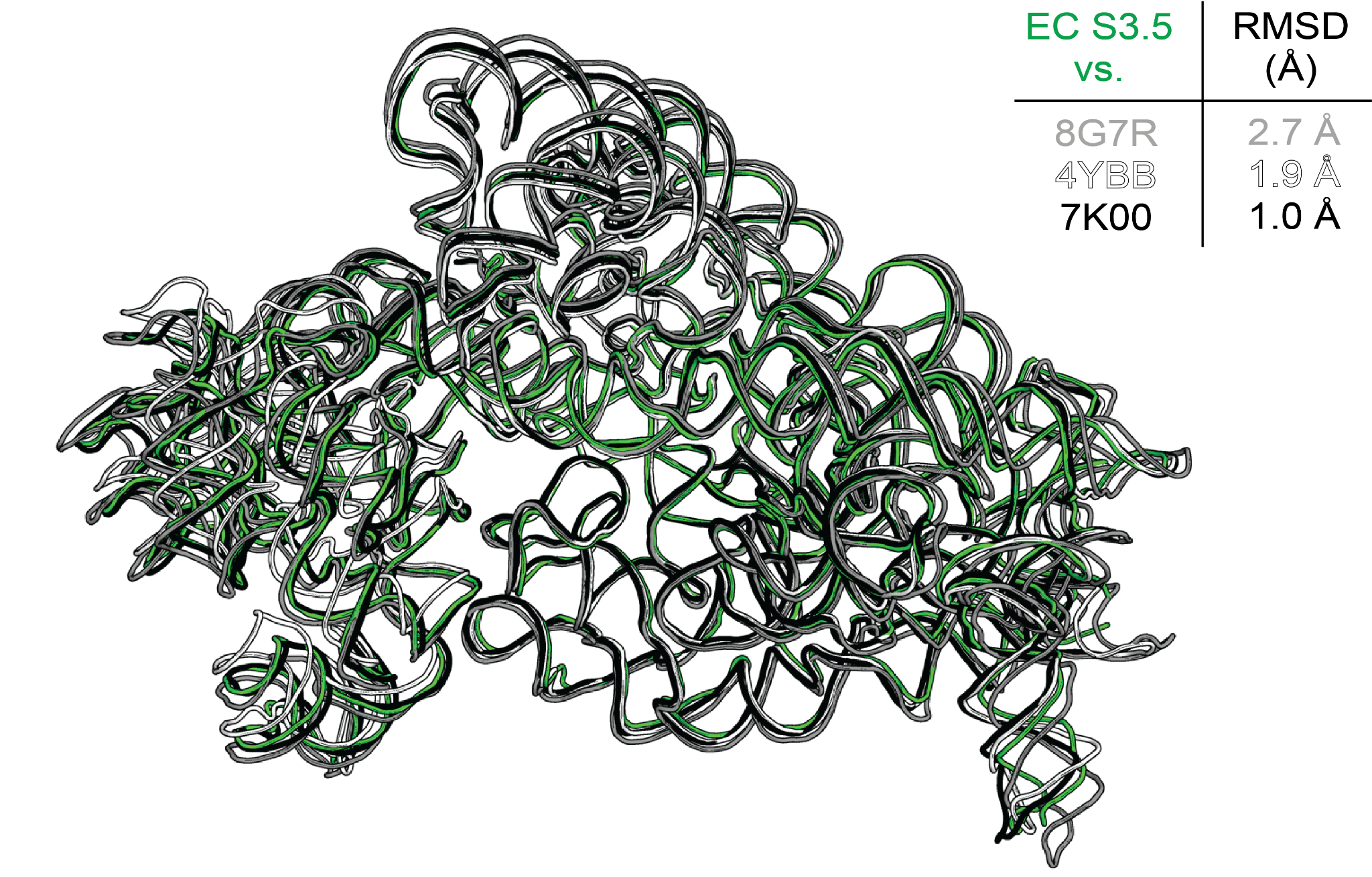

**Supplementary Figure 3. Structural comparison of EC-S3.5 with reference wildtype *E. coli* wildtype ribosomes.**

EC-S3.5 was superimposed onto the highest-resolution wildtype *E. coli* structures (PDB 7K00, 4YBB, 8G7R) to evaluate the global structural differences. RMSD values for each pairwise comparison are shown.

#### SI Figure 4

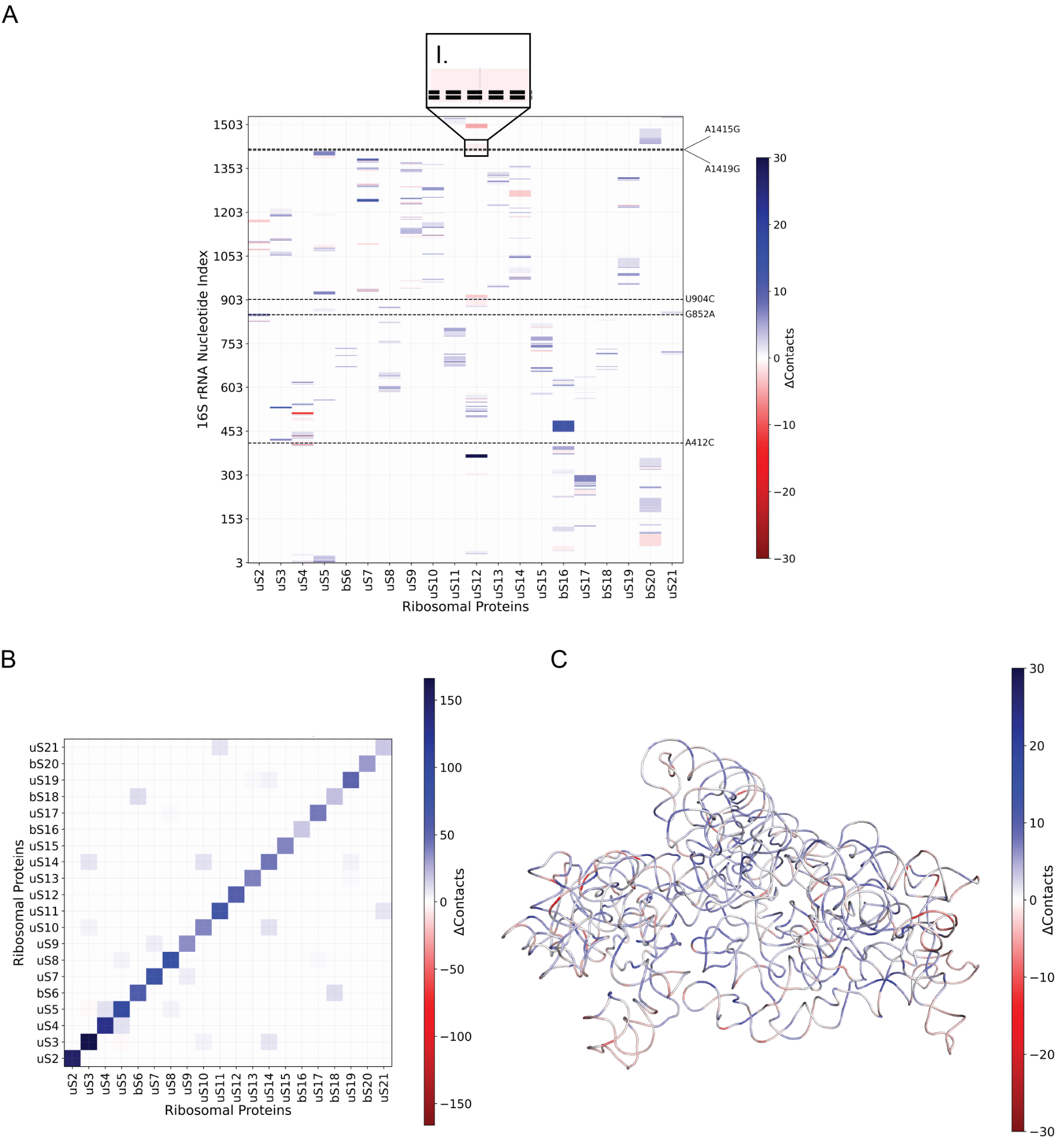

**Supplementary Figure 4. Difference in the number of pairwise contacts within 4Å between EC-S3.5 and EC-WT (PDB ID 8G7R) ribosomes.**

**A.** Differential contacts between 16S rRNA nucleotides and ribosomal proteins. Each cell depicts the net gain or loss of atomic contacts per nucleotide-protein pair, calculated by subtracting the contact counts observed in the reference structure from those in the evolved ribosome.

I. The disruption of interactions at the RNA-protein interface with uS12 due to the mutation A1415G.

**B.** Differences in contacts within and between the 30S ribosomal subunit proteins. Matrix cells represent the net change in contacts between each pair of proteins or within individual proteins (diagonal).

**C.** Differential contacts among 16S rRNA nucleotides (intra-RNA interactions). Each cell indicates the net change in contacts between nucleotide pairs across the structures compared. Colors encode the direction and magnitude of the change, with red indicating relative gains and blue indicating losses.

For all panels, contacts were defined as all interatomic distances ≤ 4 Å. Color scales are centered at zero and symmetrically scaled to the maximum absolute difference observed within each matrix. Nucleotides are indexed sequentially along the vertical and horizontal axes, and ribosomal proteins are ordered canonically from uS2 to uS21, including bacterial-specific proteins bS6, bS16, bS18, and bS20. Contact pairs were filtered to include only those where both residues or nucleotides had average Q-scores ≥ 0.4.

#### SI Figure 5

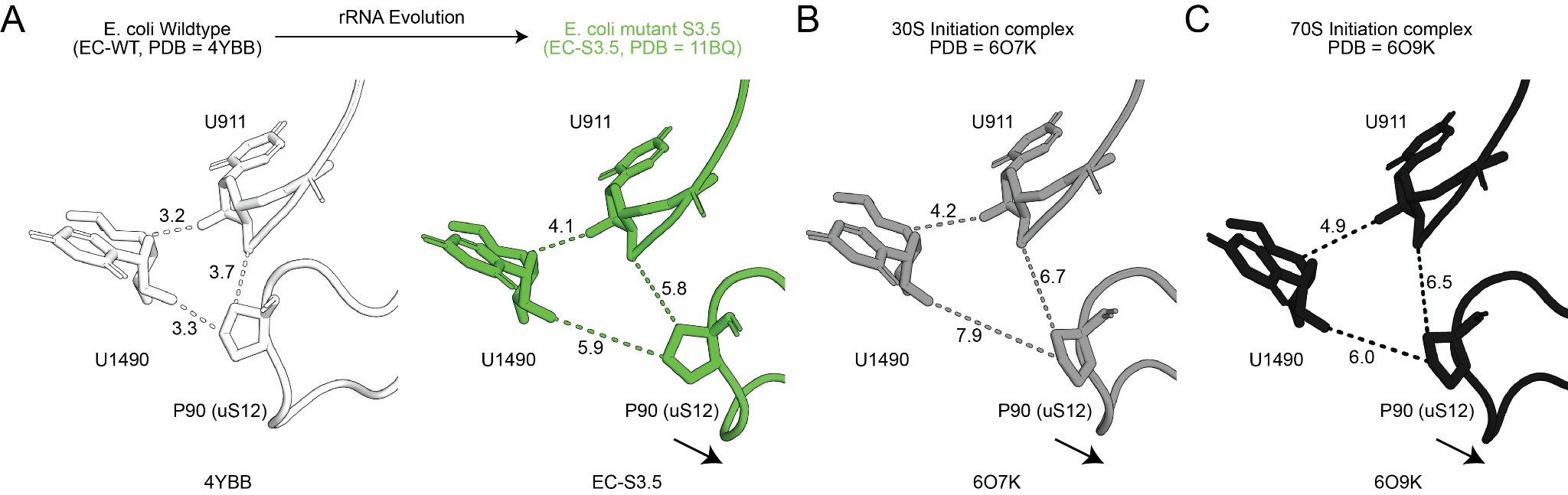

**Supplementary Figure 5. Comparison of EC-S3.5 with the initiation complexes.** To understand whether disruption of interactions at the uS12-16SrRNA interface plays a role in translation initiation, we compared our structure with A. *E.coli* wildtype (PDB ID 4YBB - white), B. the 30S initiation complex (PDB ID 6O7K - grey) and C. the 70S initiation complex (PDB ID 6O9K - black). Dashed lines represent distances and arrows represent the direction of displacement of the uS12 backbone.

#### SI Figure 6

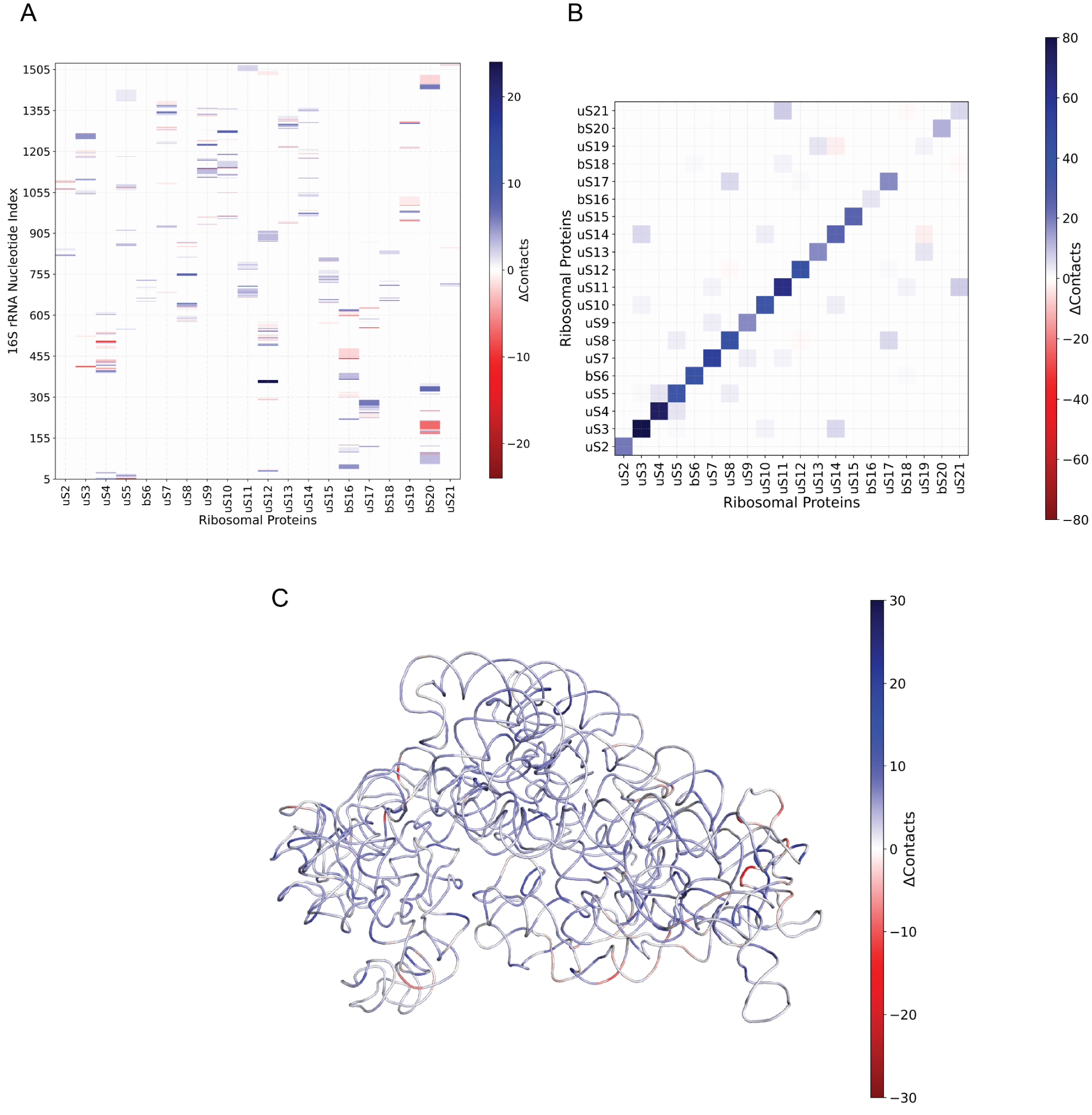

**Supplementary Figure 6. Difference in the number of pairwise contacts within 4Å between PA-ST and PA-WT (PDB ID 7UNU) ribosomes.**

**A.** Differential contacts between 16S rRNA nucleotides and ribosomal proteins. Each cell depicts the net gain or loss of atomic contacts per nucleotide-protein pair, calculated by subtracting the contact counts observed in the reference structure from those in the evolved ribosome.

**B.** Differences in contacts within and between the 30S ribosomal subunit proteins. Matrix cells represent the net change in contacts between each pair of proteins or within individual proteins (diagonal).

**C.** Differential contacts among 16S rRNA nucleotides (intra-RNA interactions). Each cell indicates the net change in contacts between nucleotide pairs across the structures compared. Colors encode the direction and magnitude of the change, with red indicating relative gains and blue indicating losses. For all panels, contacts were defined as all interatomic distances ≤4 Å. Color scales are centered at zero and symmetrically scaled to the maximum absolute difference observed within each matrix. Nucleotides are indexed sequentially along the vertical and horizontal axes, and ribosomal proteins are ordered canonically from uS2 to uS21, including bacterial-specific proteins bS6, bS16, bS18 and bS20. Contact pairs were filtered to include only those where both residues or nucleotides had average Q-scores ≥ 0.4.

#### SI Figure 7

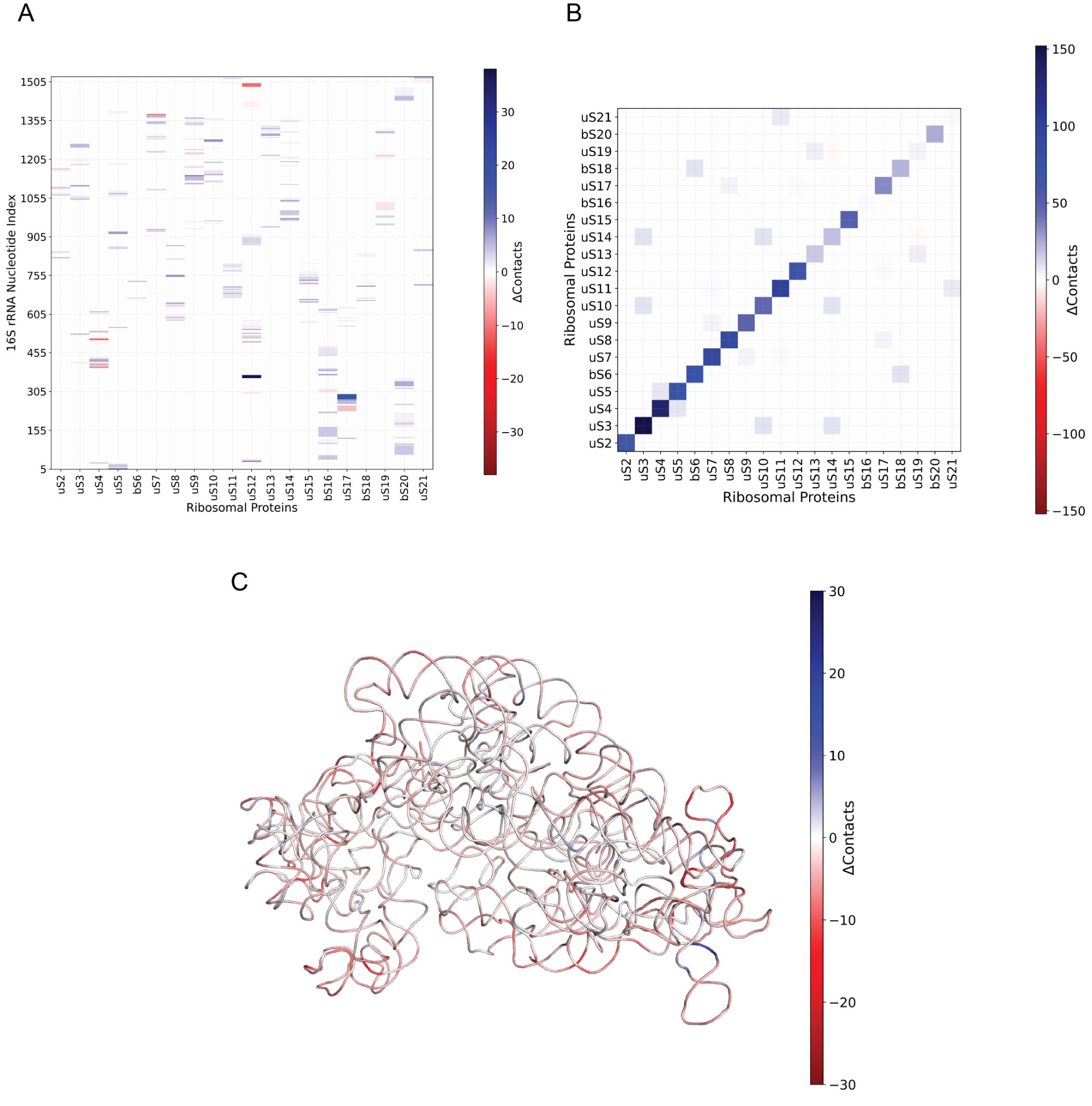

**Supplementary Figure 7. Difference in the number of pairwise contacts within 4Å between PA-ST and EC-WT (PDB ID 8G7R) ribosomes.**

**A.** Differential contacts between 16S rRNA nucleotides and ribosomal proteins. Each cell depicts the net gain or loss of atomic contacts per nucleotide-protein pair, calculated by subtracting the contact counts observed in the reference structure from those in the evolved ribosome.

**B.** Differences in contacts within and between the 30S ribosomal subunit proteins. Matrix cells represent the net change in contacts between each pair of proteins or within individual proteins (diagonal).

**C.** Differential contacts among 16S rRNA nucleotides (intra-RNA interactions). Each cell indicates the net change in contacts between nucleotide pairs across the structures compared. Colors encode the direction and magnitude of the change, with red indicating relative gains and blue indicating losses. For all panels, contacts were defined as all interatomic distances ≤4 Å. Color scales are centered at zero and symmetrically scaled to the maximum absolute difference observed within each matrix. Nucleotides are indexed sequentially along the vertical and horizontal axes, and ribosomal proteins are ordered canonically from uS2 to uS21, including bacterial-specific proteins bS6, bS16, bS18 and bS20. Contact pairs were filtered to include only those where both residues or nucleotides had average Q-scores ≥ 0.4.

#### SI Figure 8

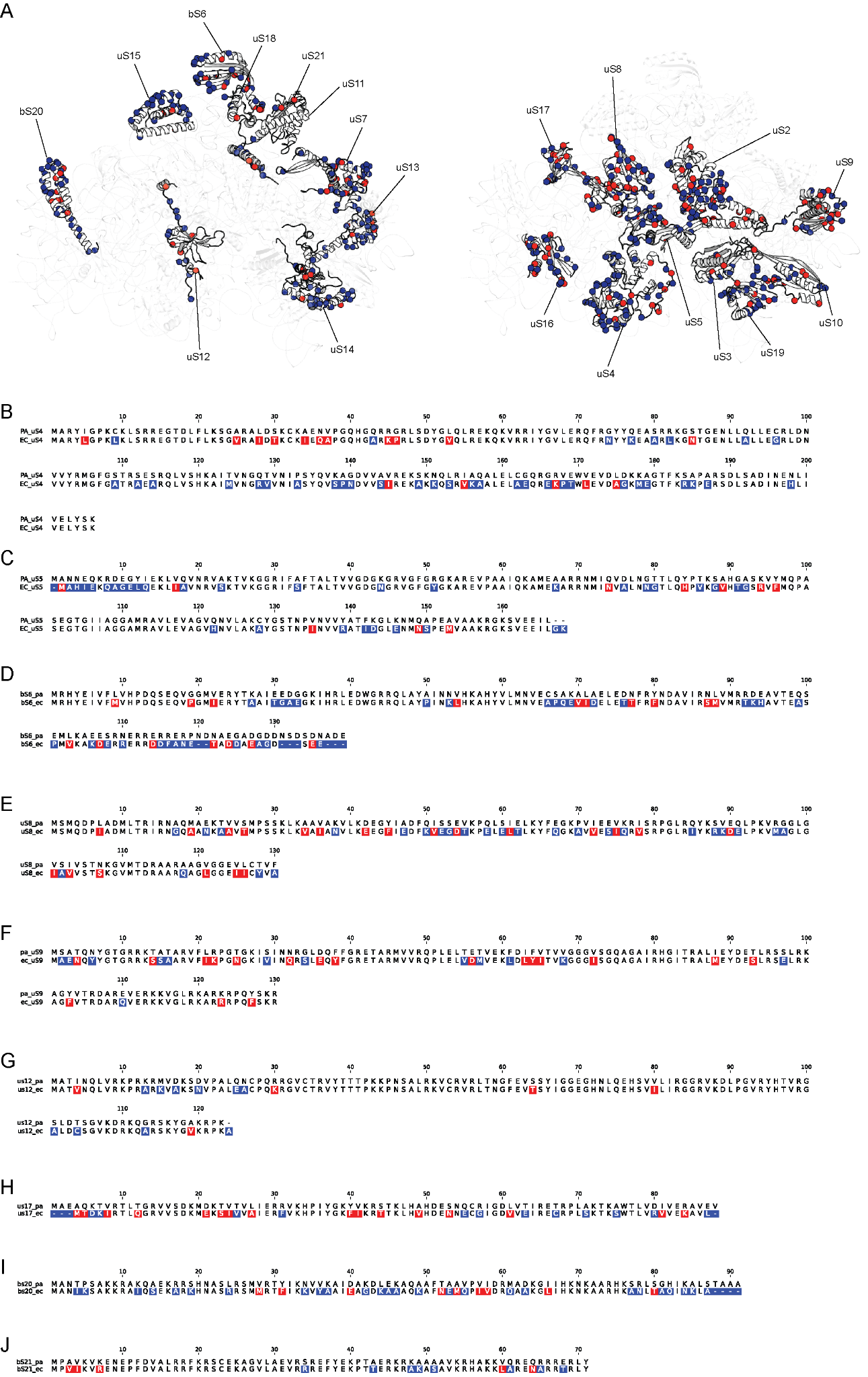

**Supplementary Figure 8. Conservative and non-conservative substitutions between *P. aeruginosa* and *E. coli* ribosomal proteins:**

**A.** The *E. coli* structure (7K00) was used to indicate the positions of conservative (red spheres) and non-conservative (blue spheres) changes between *E. coli* and *P. aeruginosa* r-proteins identified to be structurally impacted with rRNA evolution. A sequence alignment is shown for each of these r-proteins to indicate the specific residue changes: **B.** uS4, **C.** uS5, **D.** bS6, **E.** uS8, **F.** uS9, **G.** uS12, **H.** uS17, **I.** bS20, and **J.** bS21. Sequence alignments were done with cluswalW (31), and colouring of sequence alignments done with a custom Python script.

#### SI Figure 9

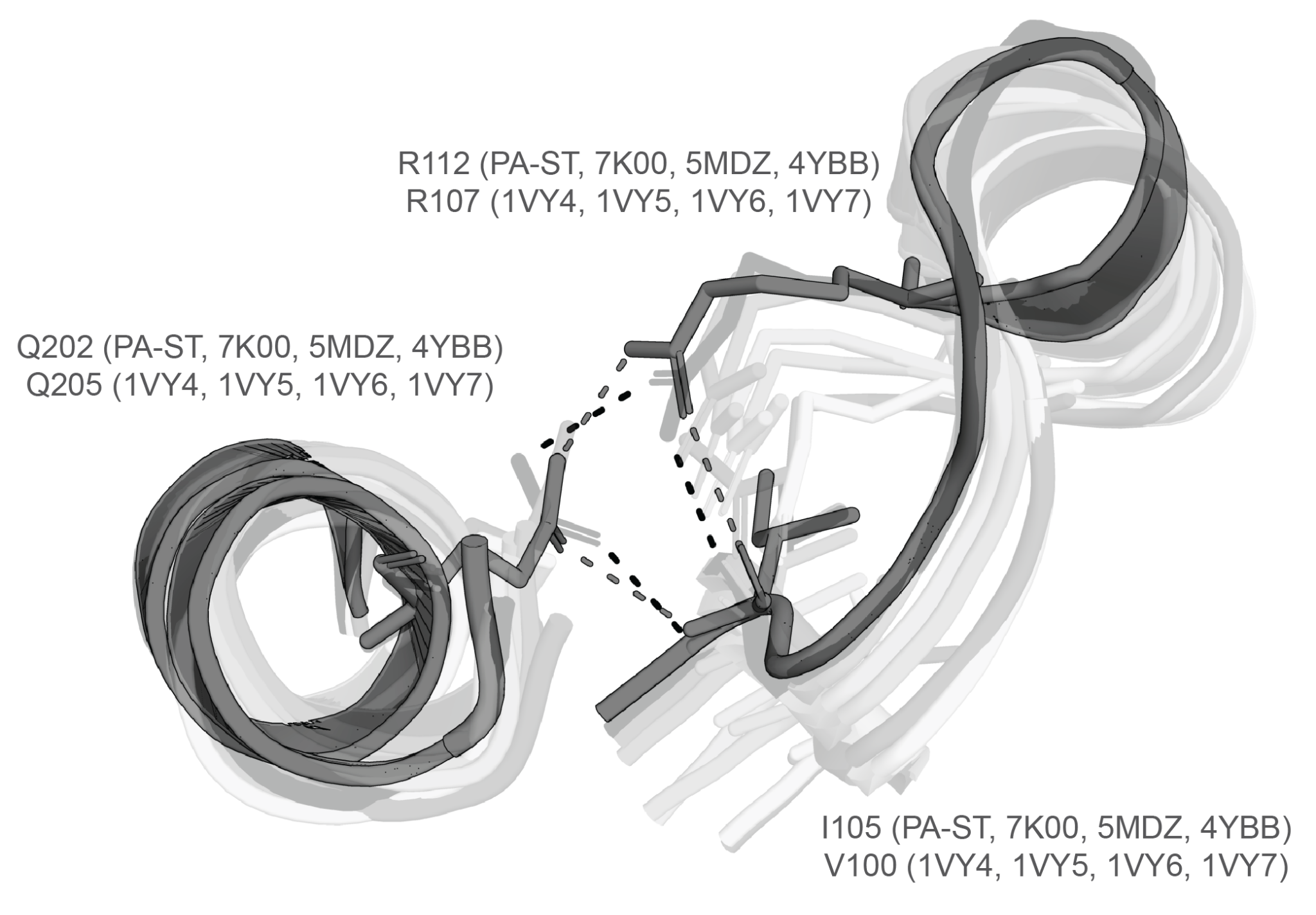

**Supplementary Figure 9. Validation of PA-ST structural changes at the uS4-uS5 interface.**

To understand the significance of the observed changes in PA-ST structure at the uS4-uS5 interface, namely a non-native hydrogen bond between uS4 I105 and uS5 E202, we looked in seven ribosome structures for the presence of this interaction (1VY4, 1VY5, 1VY6, 1VY7, 7k00, 4YBB, and 5MDZ).

#### SI Figure 10

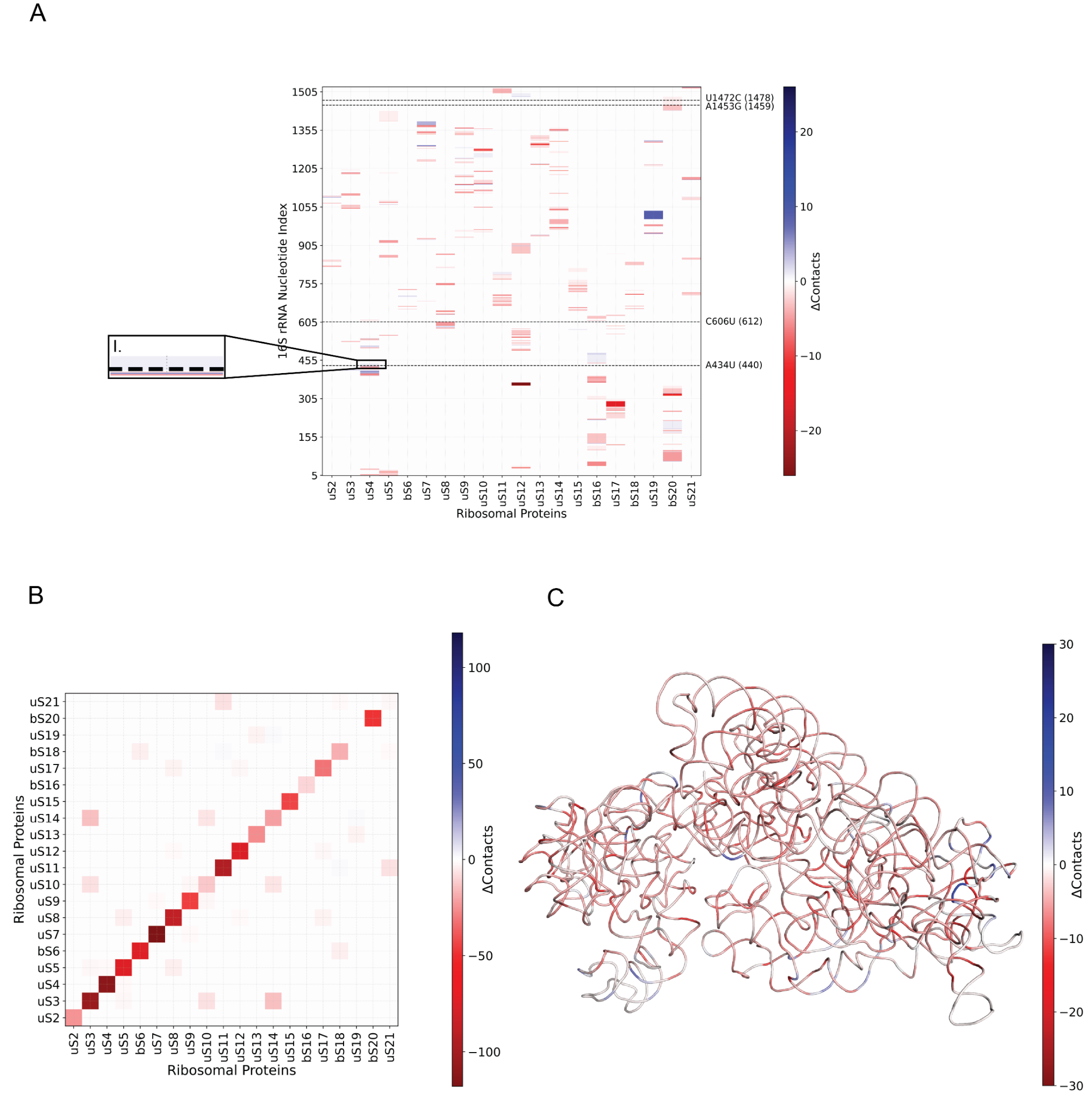

**Supplementary Figure 10. Difference in the number of pairwise contacts within 4Å between PA-S3.3 and PA-ST ribosomes.**

**A.** Differential contacts between 16S rRNA nucleotides and ribosomal proteins. Each cell depicts the net gain or loss of atomic contacts per nucleotide-protein pair, calculated by subtracting the contact counts observed in the reference structure from those in the evolved ribosome.

I. Stabilization of RNA-protein interactions with uS4 due to A434U.

**B.** Differences in contacts within and between the 30S ribosomal subunit proteins. Matrix cells represent the net change in contacts between each pair of proteins or within individual proteins (diagonal).

**C.** Differential contacts among 16S rRNA nucleotides (intra-RNA interactions). Each cell indicates the net change in contacts between nucleotide pairs across the structures compared. Colors encode the direction and magnitude of the change, with red indicating relative gains and blue indicating losses. For all panels, contacts were defined as all interatomic distances ≤4 Å. Color scales are centered at zero and symmetrically scaled to the maximum absolute difference observed within each matrix. Nucleotides are indexed sequentially along the vertical and horizontal axes, and ribosomal proteins are ordered canonically from uS2 to uS21, including bacterial-specific proteins bS6, bS16, bS18 and bS20. Dashed lines indicate positions of the mutated nucleotides. Contact pairs were filtered to include only those where both residues or nucleotides had average Q-scores ≥ 0.4.

#### SI Figure 11

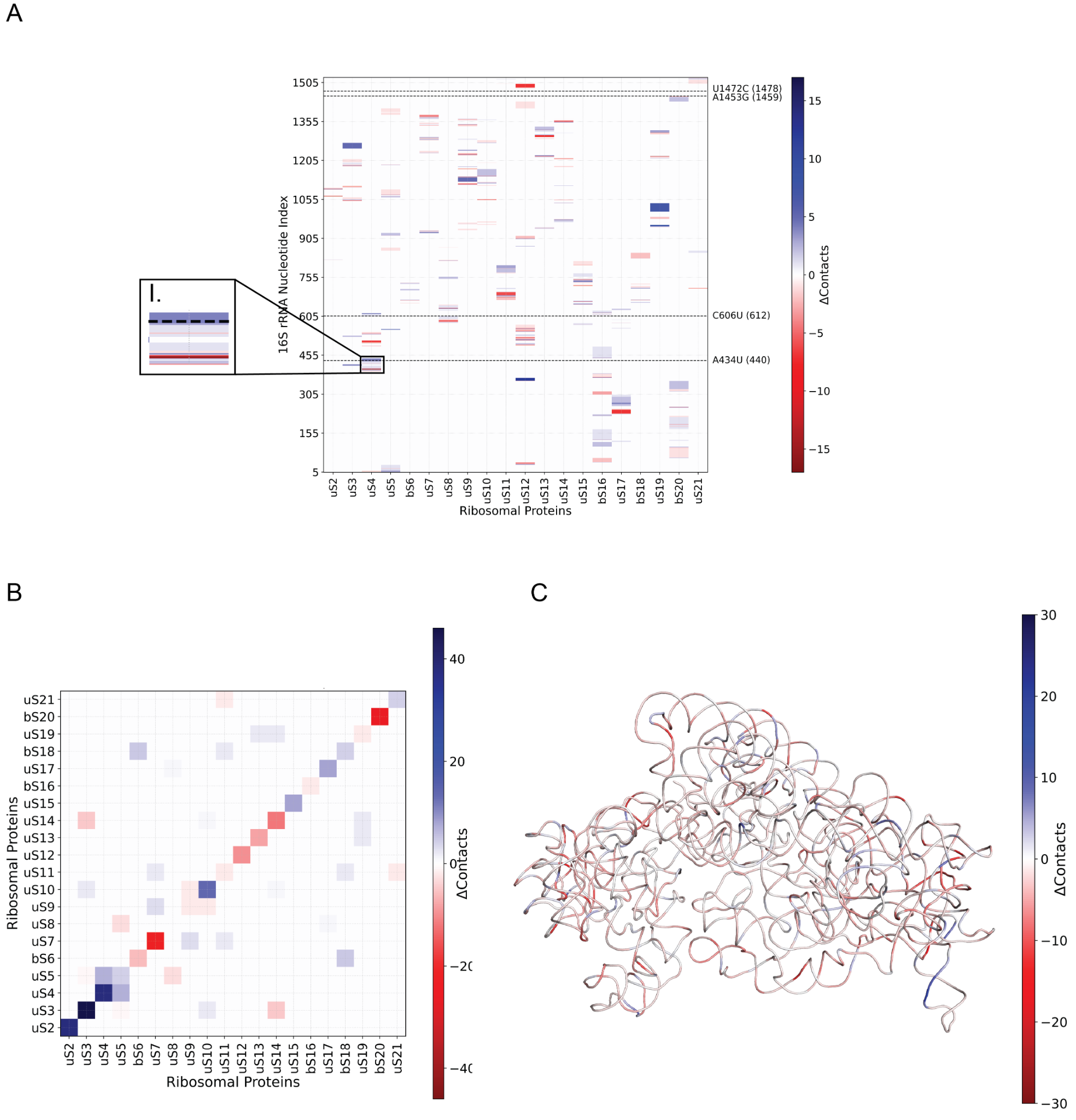

**Supplementary Figure 11. Difference in the number of pairwise contacts within 4Å between PA-S3.3 and EC-WT (PDB ID 8G7R) ribosomes.**

**A.** Differential contacts between 16S rRNA nucleotides and ribosomal proteins. Each cell depicts the net gain or loss of atomic contacts per nucleotide-protein pair, calculated by subtracting the contact counts observed in the reference structure from those in the evolved ribosome. Insets indicate the disruption or stabilization of interactions at various RNA-protein interfaces due to the mutations which includes:

I. Stabilization of uS4 due to A434U

**B.** Differences in contacts within and between the 30S ribosomal subunit proteins. Matrix cells represent the net change in contacts between each pair of proteins or within individual proteins (diagonal).

**C.** Differential contacts among 16S rRNA nucleotides (intra-RNA interactions). Each cell indicates the net change in contacts between nucleotide pairs across the structures compared. Colors encode the direction and magnitude of the change, with red indicating relative gains and blue indicating losses. For all panels, contacts were defined as all interatomic distances ≤4 Å. Color scales are centered at zero and symmetrically scaled to the maximum absolute difference observed within each matrix. Nucleotides are indexed sequentially along the vertical and horizontal axes, and ribosomal proteins are ordered canonically from uS2 to uS21, including bacterial-specific proteins bS6, bS16, bS18 and bS20. Dashed lines indicate positions of the mutated nucleotides. Contact pairs were filtered to include only those where both residues or nucleotides had average Q-scores ≥ 0.4.

#### SI Figure 12

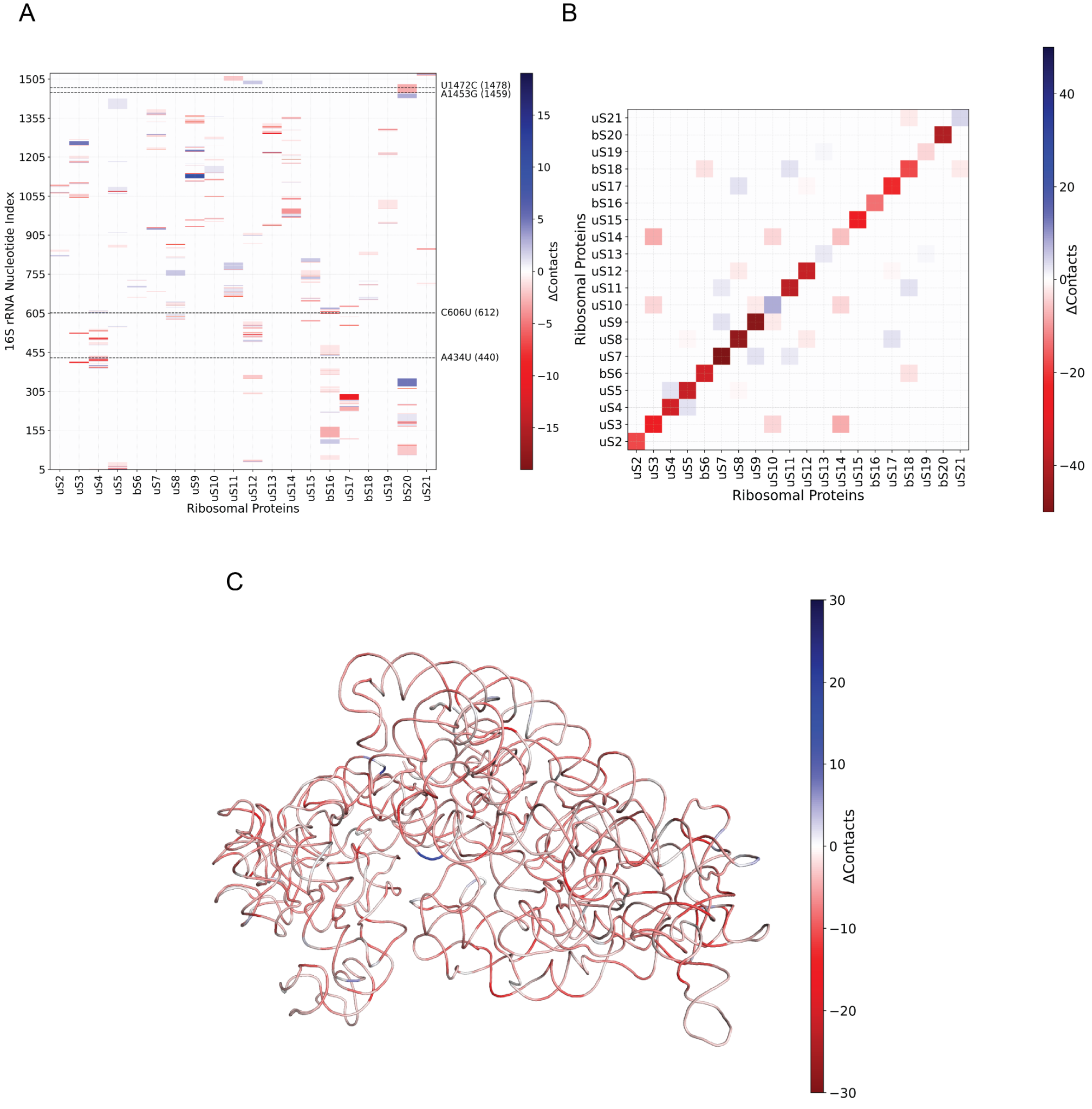

**Supplementary Figure 12. Difference in the number of pairwise contacts within 4Å between PA-S3.3 and PA-WT (PDB ID 7UNU) ribosomes.**

**A.** Differential contacts between 16S rRNA nucleotides and ribosomal proteins. Each cell depicts the net gain or loss of atomic contacts per nucleotide-protein pair, calculated by subtracting the contact counts observed in the reference structure from those in the evolved ribosome.

**B.** Differences in contacts within and between the 30S ribosomal subunit proteins. Matrix cells represent the net change in contacts between each pair of proteins or within individual proteins (diagonal).

**C.** Differential contacts among 16S rRNA nucleotides (intra-RNA interactions). Each cell indicates the net change in contacts between nucleotide pairs across the structures compared. Colors encode the direction and magnitude of the change, with red indicating relative gains and blue indicating losses. For all panels, contacts were defined as all interatomic distances ≤4 Å. Color scales are centered at zero and symmetrically scaled to the maximum absolute difference observed within each matrix. Nucleotides are indexed sequentially along the vertical and horizontal axes, and ribosomal proteins are ordered canonically from uS2 to uS21, including bacterial-specific proteins bS6, bS16, bS18 and bS20. Dashed lines indicate positions of the mutated nucleotides. Contact pairs were filtered to include only those where both residues or nucleotides had average Q-scores ≥ 0.4.

#### SI Figure 13

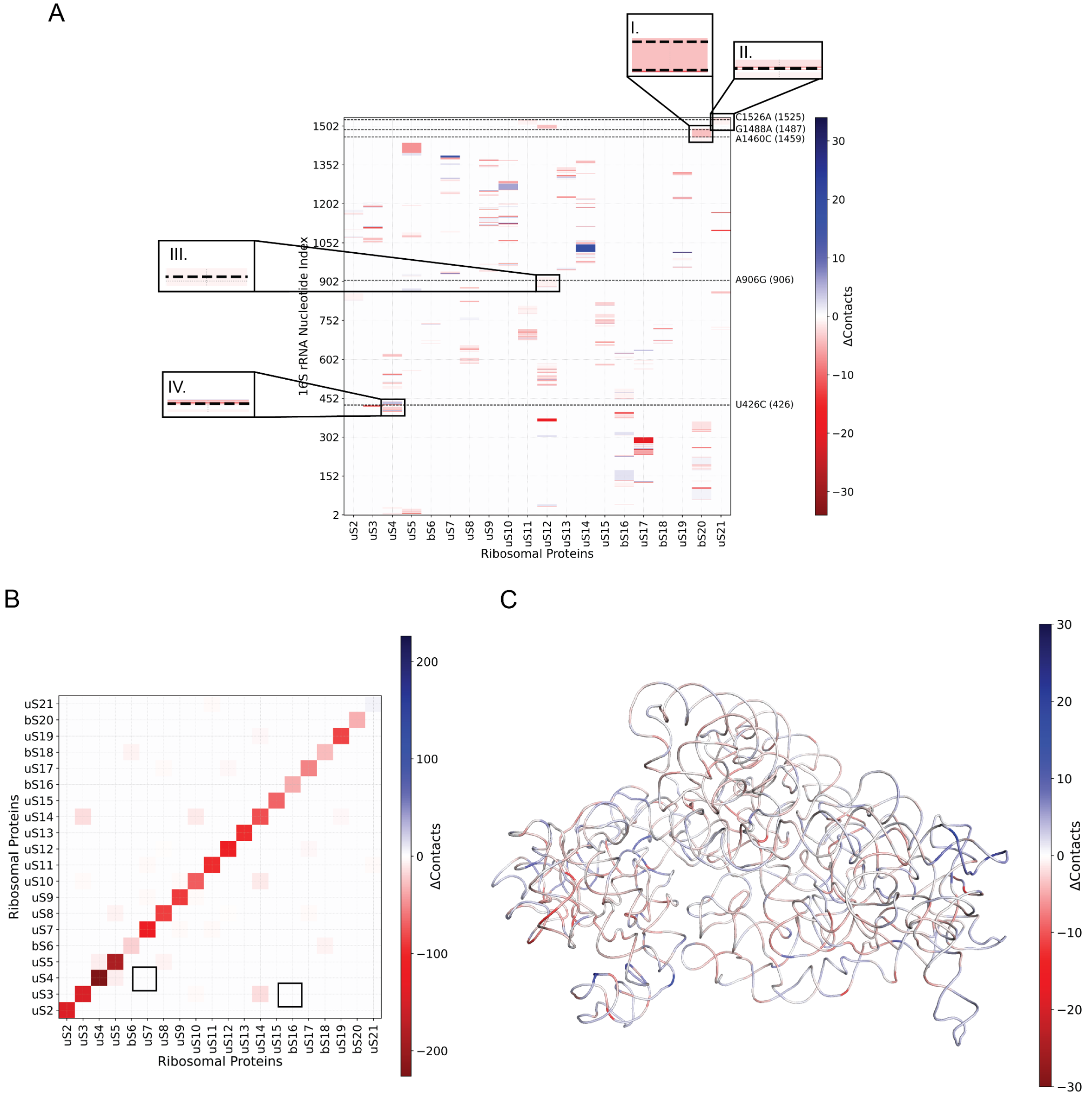

**Supplementary Figure 13. Difference in the number of pairwise contacts within 4Å between VC-S4.4 and VC-ST ribosomes.**

**A.** Differential contacts between 16S rRNA nucleotides and ribosomal proteins. Each cell depicts the net gain or loss of atomic contacts per nucleotide-protein pair, calculated by subtracting the contact counts observed in the reference structure from those in the evolved ribosome. Insets indicate the disruption of interactions at various RNA-protein interfaces due to the mutations which includes:

I. bS20 as a result of G1488A
II. uS21 due to C1526A
III. uS12 due to A906G and
IV. uS4 due to U426C

**B.** Differences in contacts within and between the 30S ribosomal subunit proteins. Matrix cells represent the net change in contacts between each pair of proteins or within individual proteins (diagonal).

**C.** Differential contacts among 16S rRNA nucleotides (intra-RNA interactions). Each cell indicates the net change in contacts between nucleotide pairs across the structures compared. Colors encode the direction and magnitude of the change, with red indicating relative gains and blue indicating losses. For all panels, contacts were defined as all interatomic distances ≤4 Å. Color scales are centered at zero and symmetrically scaled to the maximum absolute difference observed within each matrix. Nucleotides are indexed sequentially along the vertical and horizontal axes, and ribosomal proteins are ordered canonically from uS2 to uS21, including bacterial-specific proteins bS6, bS16, bS18, and bS20. Dashed lines indicate positions of the mutated nucleotides. Contact pairs were filtered to include only those where both residues or nucleotides had average Q-scores ≥ 0.4.

#### SI Figure 14

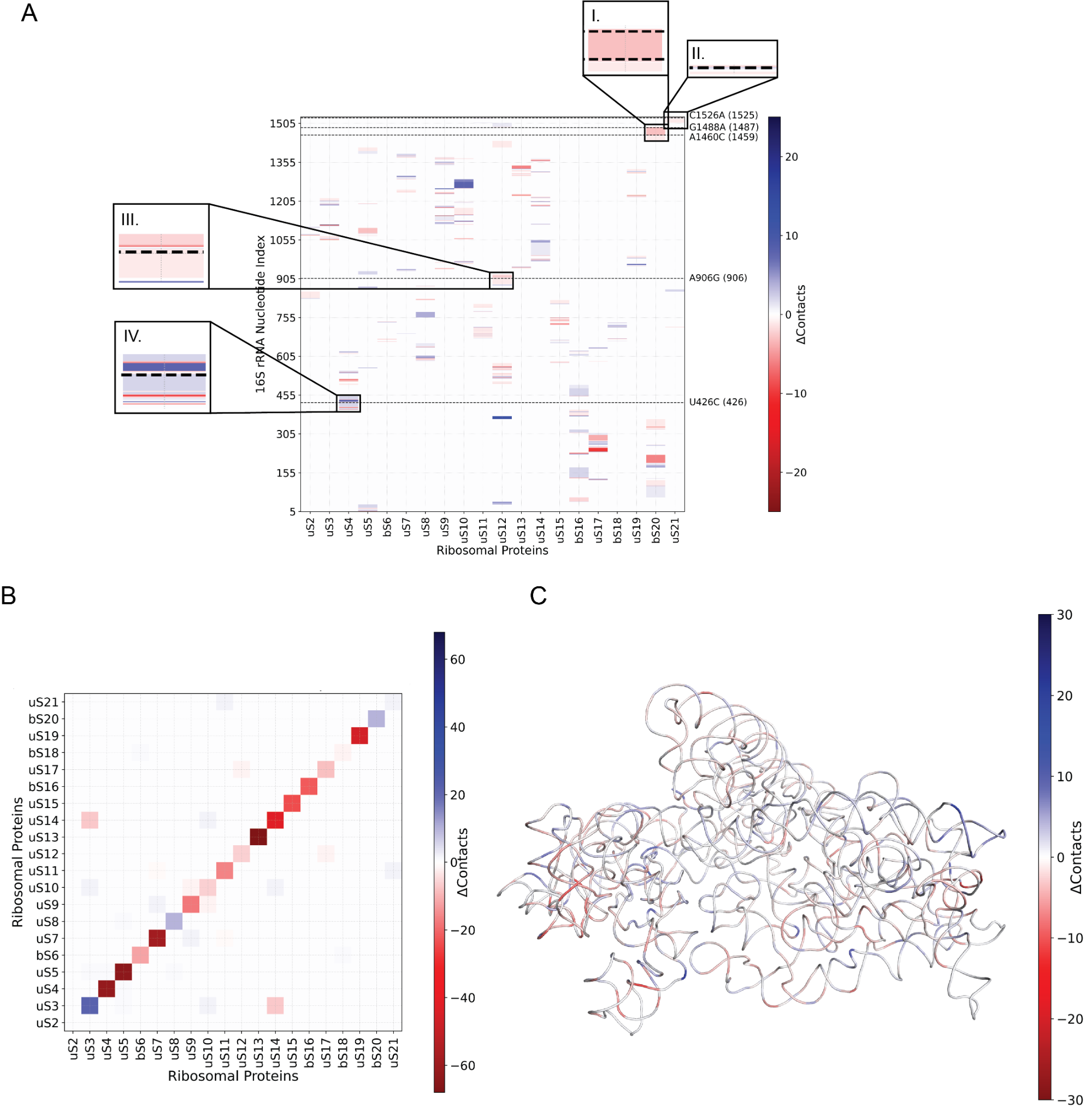

**Supplementary Figure 14. Difference in the number of pairwise contacts within 4Å between VC-S4.4 and EC-WT (PDB ID 8G7R) ribosomes.**

**A.** Differential contacts between 16S rRNA nucleotides and ribosomal proteins. Each cell depicts the net gain or loss of atomic contacts per nucleotide-protein pair, calculated by subtracting the contact counts observed in the reference structure from those in the evolved ribosome. Insets indicate the disruption of interactions at various RNA-protein interfaces due to the mutations which includes:

I. bS20 as a result of G1488A
II. uS21 due to C1526A
III. uS12 due to A906G and
IV. uS4 due to U426C

**B.** Differences in contacts within and between the 30S ribosomal subunit proteins. Matrix cells represent the net change in contacts between each pair of proteins or within individual proteins (diagonal).

**C.** Differential contacts among 16S rRNA nucleotides (intra-RNA interactions). Each cell indicates the net change in contacts between nucleotide pairs across the structures compared. Colors encode the direction and magnitude of the change, with red indicating relative gains and blue indicating losses. For all panels, contacts were defined as all interatomic distances ≤4 Å. Color scales are centered at zero and symmetrically scaled to the maximum absolute difference observed within each matrix. Nucleotides are indexed sequentially along the vertical and horizontal axes, and ribosomal proteins are ordered canonically from uS2 to uS21, including bacterial-specific proteins bS6, bS16, bS18 and bS20. Dashed lines indicate positions of the mutated nucleotides. Contact pairs were filtered to include only those where both residues or nucleotides had average Q-scores ≥ 0.4.

#### SI Figure 15

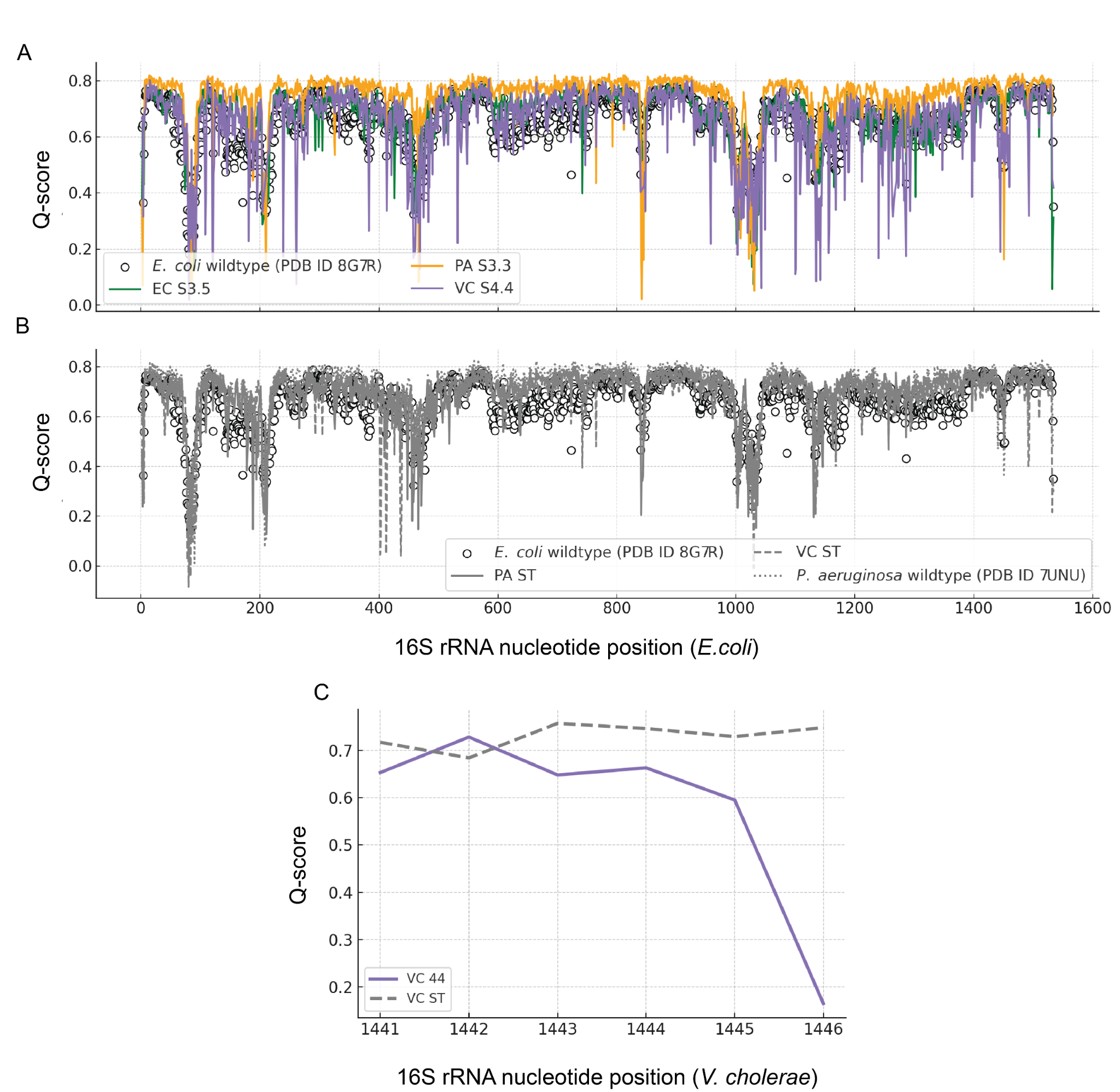

**Supplementary Figure 15. Q-score analysis for 16S rRNA:**

**A.** Plots represent Q-scores for VC-S4.4 evo (violet), PA-S3.3 (orange), EC-S3.5 (green), and

**B.** Q-scores for PA-ST (solid line), PA-WT (PDB ID 7UNU - dotted line), VC-ST(dashed line) and *E. coli* wildtype (hollow circles with black border) (PDB ID 8G7R) ribosomes.

**C.** For nucleotides 1441-1446 in VC-S4.4 (violet) and VC-ST (dashed line)

*Inset: Q-score for nucleotides 1442-1454 of h44. The Q-scores are obtained using the “Model-map-Q-score” plugin in chimeraX (68),(69) .

#### SI Figure 16

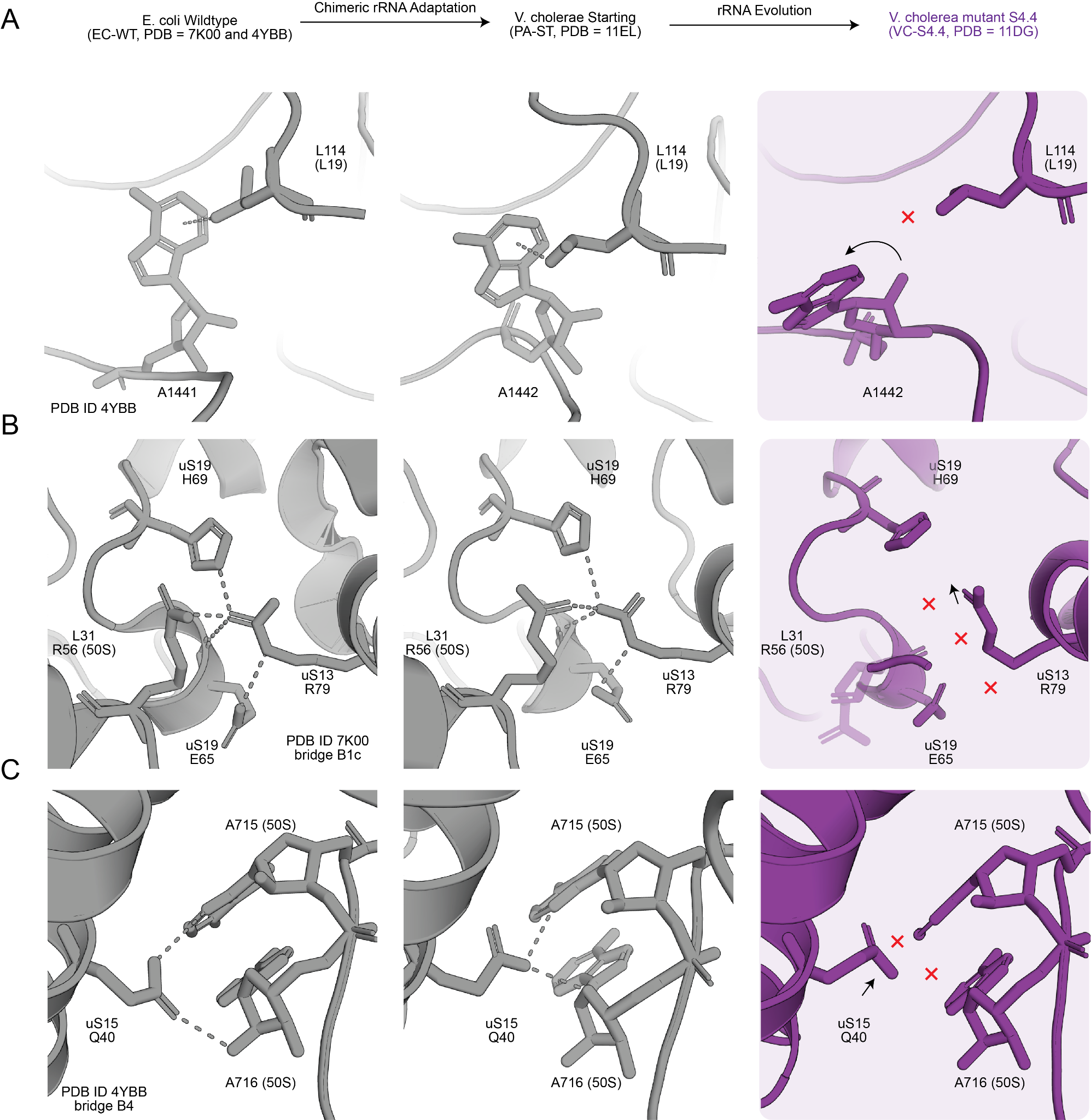

**Supplementary Figure 16. Disruption of specific intersubunit interactions in VC-S4.4**

Structural comparison of selected 30S-50S interactions in VC-S4.4 with EC-WT (PDB IDs 7K00 and 4YBB) and VC-ST. Red exes indicate contacts lost in VC-S4.4 relative to both EC-WT and VC-ST. Black arrows indicate the direction of displacement or conformational change.

**A.** CH-π interaction between A1442 of 16S rRNA (A1441 in E. coli) and L114 of 50S protein L19.

**B.** B1c bridge involving E65 and H69 of uS19, R79 of uS13, and R56 of 50S protein L31. In VC-S4.4, the density for the L31 R56 side chain is sparse; it is therefore shown as semi-transparent sticks to indicate side chain disorder. PDB ID 7K00 was used as the EC-WT reference for this comparison because residues R56-I66 are not modeled in 4YBB or 8G7R.

**C.** B4 interaction between Q40 of uS15 and A715/A716 of 23S rRNA in the 50S subunit.

#### SI Figure 17

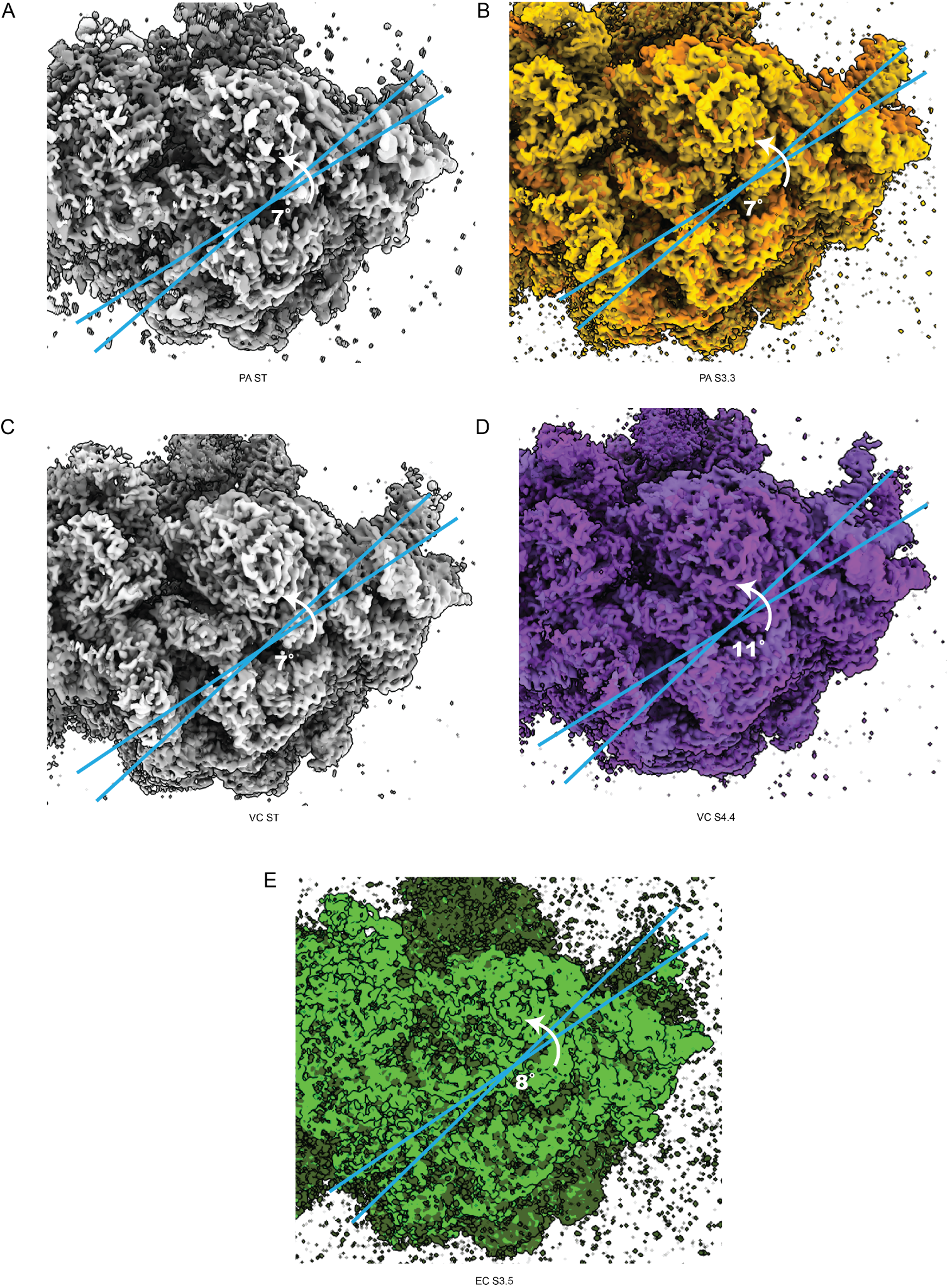

**Supplementary Figure 17.** **Conformational heterogeneity in chimeric and non chimeric ribosomes:** Volumes obtained from 3D classification in cryoSPARC(23) using **A. PA-ST, B. PA S3.3, C. VC-ST, D. VC S4.4 and E. EC S3.5** datasets are shown. Volumes showing the largest variability in terms of the rotation angle of the 30S are displayed. Both the volumes in each of the panels are contoured at the same level (3.6 𝛔). The arrow indicates the direction of ratcheting motion. The volumes were superposed using the “fit in Map” module in ChimeraX(69).
